# The cortical microenvironment drives early immune organization and controls early osteoclastogenesis in bone healing

**DOI:** 10.1101/2025.11.03.684298

**Authors:** Anne Noom, Hülya Z. Oktay, Duncan M. Morgan, Sandy Kroh, Agnes Ellinghaus, Merle Kochan, Olufemi Bolaji, Ralf Uecker, Robert Günther, Christian H. Bucher, Antigoni Triantafyllopoulou, Simon Haas, Katharina Schmidt-Bleek, Anja E. Hauser, Birgit Sawitzki, Georg N. Duda

**Affiliations:** Julius Wolff Institute (JWI), Berlin Institute of Health at Charité – Universitätsmedizin Berlin, 13353 Berlin, Germany; BIH Center for Regenerative Therapies (BCRT), Berlin Institute of Health at Charité – Universitätsmedizin Berlin, 13353 Berlin, Germany; Center for Musculoskeletal Surgery, Charité – Universitätsmedizin Berlin, 13353 Berlin, Germany; Department of Infectious Diseases and Respiratory Medicine, Charité – Universitätsmedizin Berlin, Corporate Member of Freie Universität Berlin and Humboldt University of Berlin, 13353 Berlin, Germany; Center of Immunomics, Berlin Institute of Health at Charité – Universitätsmedizin Berlin, 13353 Berlin, Germany; Wyss Institute for Biologically Inspired Engineering and John A. Paulson School of Engineering and Applied Sciences, Harvard University, Cambridge, MA 02138, USA; Department of Rheumatology and Clinical Immunology, Charité-Universitätsmedizin Berlin Corporate Member of Freie Universität Berlin and Humboldt-Universität zu Berlin, 10117 Berlin, Germany; Immune Dynamics, Deutsches Rheuma-Forschungszentrum (DRFZ), a Leibniz Institute, 10117 Berlin, Germany; Institute of Microbiology, Infectious Diseases and Immunology (I-MIDI), Charité-Universitätsmedizin Berlin Campus Benjamin Franklin, 12203 Berlin, Germany; Deutsches Rheuma-Forschungszentrum (DRFZ), a Leibniz Institute, 10117 Berlin, Germany; Berlin Institute of Health at Charité – Universitätsmedizin Berlin, 10115 Berlin, Germany; Berlin Institute of Medical Systems Biology, Max Delbrück Center for Molecular Medicine in the Helmholtz Association, 10115 Berlin, Germany; Precision Healthcare University Research Institute, Queen Mary University of London, London, United Kingdom

**Author notes:** Correspondence: Prof. Dr.-Ing. Georg N. Duda, Julius Wolff Institute and Berlin Institute of Health Center for Regenerative Therapies, Berlin Institute of Health at Charité – Universitätsmedizin Berlin, Augustenburger Platz 1, D-13353 Berlin, Germany.

## Abstract

Bone regeneration is a complex, tightly-regulated process involving coordinated interactions of immune and stromal cells. Early phases of healing rely on the timely clearance of debris, a task primarily carried out by macrophages and osteoclasts. However, the sequence of events leading to the presence of osteoclasts at the fracture site and how this is shaped by local tissue microenvironments remains poorly understood, particularly at single-cell and spatial resolution. Using single-cell RNA sequencing and multi-epitope ligand cartography, we mapped the spatial organization of distinct cell compartments engaged in early fracture healing in both young and aged mice at the start of healing. Surprisingly, we found that young mice exhibited an increased presence of activated osteoclasts at day 7, concentrated within the cortical niche. This compartment was also characterized by a spatially restricted immune response with a selective accumulation of distinct macrophage types jointly interacting with neutrophils and stromal cells. This raised the possibility that local cell organization influences osteoclast precursor differentiation. We identified a distinct *Spp1*^hi^ macrophage subset restricted to the cortex, which acted as a transitional precursor population giving rise to osteoclasts. Neutrophils preceded this *Spp1*^hi^ macrophage accumulation and may promote their recruitment through chemotactic signaling. This coordination was less pronounced in aged mice despite preserved transcriptional states. In parallel, stromal cells in young animals displayed higher expression of essential niche factors further supporting local osteoclastogenesis at the cortex. Together, our findings identify distinct macrophage precursors and reveal early, cortex-specific niche activity supporting osteoclastogenesis. This provides a new framework for understanding the initiation of spatial immune–stromal interactions for the early stages of regeneration.

## Introduction

Bone fractures are frequent and affect both young and old patients. In almost all cases, fracture healing proceeds without complications to a scar-free regeneration^1–3^. Current treatments, including surgical stabilization and biological augmentation, generally support this natural healing process, but complications can still arise, causing delayed healing or even non-unions^4–6^. A better understanding of the cellular mechanisms that regulate bone regeneration is therefore still essential to improve therapeutic strategies.

Bone regeneration is a complex, highly dynamic and tightly regulated process, involving various immune cells^7,8^, such as neutrophils^9,10^ and macrophages^11,12^, osteoclasts^13,14^ and tissue forming cells such as stroma cells and osteoblasts^15^. Following fracture, the area is initially filled with a blood clot, where coagulation and platelet activation create a provisional matrix. Neutrophils are among the first immune cells to enter this environment, initiating the acute inflammatory response and releasing chemotactic signals to recruit other cell types^9,16,17^. Macrophages soon follow and play multifaceted roles throughout the healing process. Traditionally described along an M1/M2 axis, macrophages range from pro-inflammatory cells that drive early immune activation to anti-inflammatory populations that support tissue repair^11,12,18^. Within the bone environment, this functional diversity is further expanded by the presence of both monocyte-derived macrophages and long-lived tissue-resident populations such as osteal macrophages (osteomacs), which support osteoblast function and matrix homeostasis^19,20^. Their plasticity and responsiveness to local cues position macrophages as central regulators of the early healing environment.

In addition to their regulatory roles, cells of the macrophage lineage serve as precursors to a specific cell type associated with bone turnover, the osteoclasts, which are multinucleated cells specialized in resorbing mineralized bones. Osteoclast differentiation is driven by macrophage colony-stimulating factor (M-CSF; *Csf1*) and receptor activator of nuclear factor κB ligand (RANKL) signaling and is classically associated with late-stage bone healing and remodeling^21,22^. However, early histological evidence showed that osteoclastic activity begins sooner than previously assumed and increases progressively in the endosteal and cortical regions^13^. Osteoclast numbers are usually rather low but increase rapidly following fracture, accumulating at the injury site to degrade necrotic bone debris^14^. This process creates the spatial and structural conditions necessary to initiate a regenerative niche for new bone formation jointly with osteoblasts^13,23^. In such niche constellations, the concept of skeletal coupling can be found with bone resorption and formation being closely linked.

The early presence of osteoclast activity suggests that osteoclast lineage commitment may be initiated shortly after injury, rather than being restricted to the late remodeling phase. However, despite the recognized lineage potential of myeloid cells, it remains unknown which cells give rise to osteoclasts during this early bone regeneration, and how their differentiation is shaped by local tissue context *in vivo*. The process of osteoclast differentiation has been typically studied under uniform *in vitro* conditions^24^, while bone healing takes place *in vivo* in spatially distinct compartments, such as bone marrow and cortical bone including the endosteum and periosteum, that differ in cell composition^25^, signaling environment^26^ and mechanical context^27^ and may uniquely influence macrophage and osteoclast precursor behavior and cell fate. In aged individuals, impaired healing is associated with altered innate and adaptive immune responses^12^ and in parallel with an increased osteoclast activity^28^. Yet, the link between aging and macrophage versus osteoclast differentiation remains unclear. Clarifying how niche context drives macrophage and osteoclast phenotypes in early phases of bone regeneration and how this is impacted by age is critical for understanding the very early immune regulation in bone healing.

In this study, we investigated how early osteoclastogenesis is shaped by spatially organized immune-stromal interactions during bone healing. Using single-cell transcriptomics and multi-epitope ligand cartography (MELC), we identify a transitional *Spp1^hi^* macrophage subset that emerges within the cortical microenvironment and gives rise to osteoclasts in a spatially confined manner. This early differentiation trajectory is modulated by neutrophil-macrophage crosstalk and stromal-derived niche signals, which are dampened in aged mice. These findings uncover a spatially coordinated mechanism of osteoclastogenesis during regeneration and highlights the cortical microenvironment as a central regulator of early bone regeneration.

## Results

### Osteoclast activity is delayed during early fracture healing of aged mice

To better understand how tissue microenvironments and aging influence macrophage and osteoclast differentiation during bone healing, we employed a standardized osteotomy model in young (12-week-old) and aged (52-week-old) mice, as previously published^7,11,29^. In short, a 0.7 mm osteotomy was introduced in the left femur and stabilized using a unilateral external fixator (MouseExFix, RISystem, Davos, Switzerland). This model allows us to examine the native influence of tissue microenvironment on fracture healing and how this is altered with age, in the absence of any therapeutic intervention.

To determine whether osteoclasts are also active during the very early stages of bone healing, we focused on day 3 and day 7 post-osteotomy, preceding the classical remodeling phase typically assessed weeks after fracture^13^. Histological analysis of the fracture revealed the presence of bone fragments near the cortical edges at day 3, with partial clearance by day 7 (Suppl. Figure 1A-D). To functionally evaluate osteoclast activity, we performed TRAP staining at both timepoints. In young mice, we observed a clear increase in TRAP-positive cells by day 7, particularly along the cortical bone surface and around residual bone debris (Figure 1A, arrowheads), whereas this induction appeared less pronounced in aged animals (Figure 1B). This early evidence of osteoclast activity in young mice indicates that osteoclast precursors are already present at the onset of healing, complementing other macrophage types that clear the hematoma and soft tissue remnants after fracture. Given the importance of removing necrotic or damaged tissues and even bone fragments as prerequisite of any subsequent tissue regeneration^13^, these findings suggest that early osteoclast activity and their distinct differentiation from precursor cells may be a critical process for an effective fracture repair process and be altered in aged settings.

**Figure 1:**
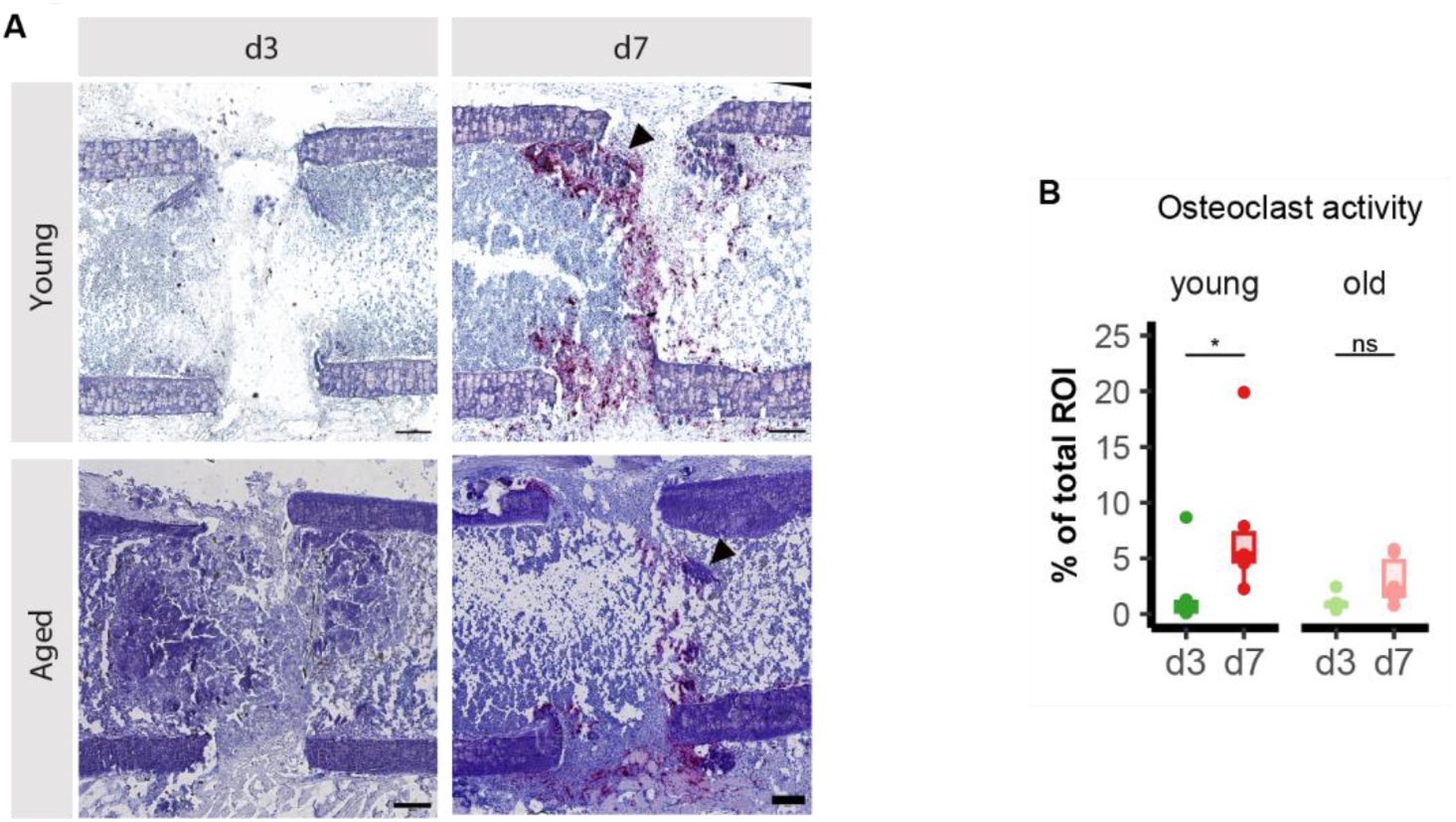
Early detection of osteoclast activity during fracture healing using TRAP staining. (A) Representative histological image showing TRAP (tartrate-resistant acid phosphatase) staining of fracture gap at day 3 and day 7 post-osteotomy of young (12-week-old) and aged (52-week-old) animals. TRAP-positive osteoclasts (red) are primarily located along the cortical bone surface (purple), reflecting localized bone resorption activity in the early phase of healing. All femurs were oriented consistently, with the proximal end on the left and the distal end on the right side of each image. Scale bar = 200 µm. (B) Quantification of TRAP-positive area within the region of interest (ROI) (0.7 mm fracture gap, including 0.4 mm proximal and distal bone segments), performed using an in-house macro in Fiji. n = 5-6 biological replicates per group; *p < 0.05, Wilcoxon test.

### Fracture-induced immune responses are highly compartmentalized

Given the enhanced osteoclast activity observed by day 7, we next sought to understand the cellular and molecular mechanisms underlying this early response. Since TRAP-positive cells were predominantly localized along the cortical bone surface, we hypothesized that distinct microenvironmental cues within the marrow and cortical compartments may differentially regulate osteoclast precursor differentiation.

To test this hypothesis, we employed single-cell RNA sequencing for young and aged mice to comprehensively profile osteoclast precursors and their local microenvironments within the marrow and cortical compartments during early fracture healing. To focus on the cell populations most relevant to bone regeneration, we analyzed cells that were recovered exclusively between the proximal and distal inner fixation pins. A two-step approach was used to separate the compartments (Figure 2A). First, bone marrow-resident cells were collected through a direct flushing of the bone cavity. Then, cortex-adherent cells were liberated by subjecting the remaining cortical fragments to enzymatic digestion using a combination of collagenase and dispase. We analyzed cells from 30 mice present at day 3 and day 7 post-osteotomy, while intact marrow and cortex of age-matched mice served as controls. As an additional control, we also analyzed cells obtained from the contralateral, uninjured femur. This approach, employed in prior studies of bone microenvironments^7,25^, allowed us to create for the first time a single-cell atlas of both the marrow and cortex during the early onset of fracture healing.

**Figure 2:**
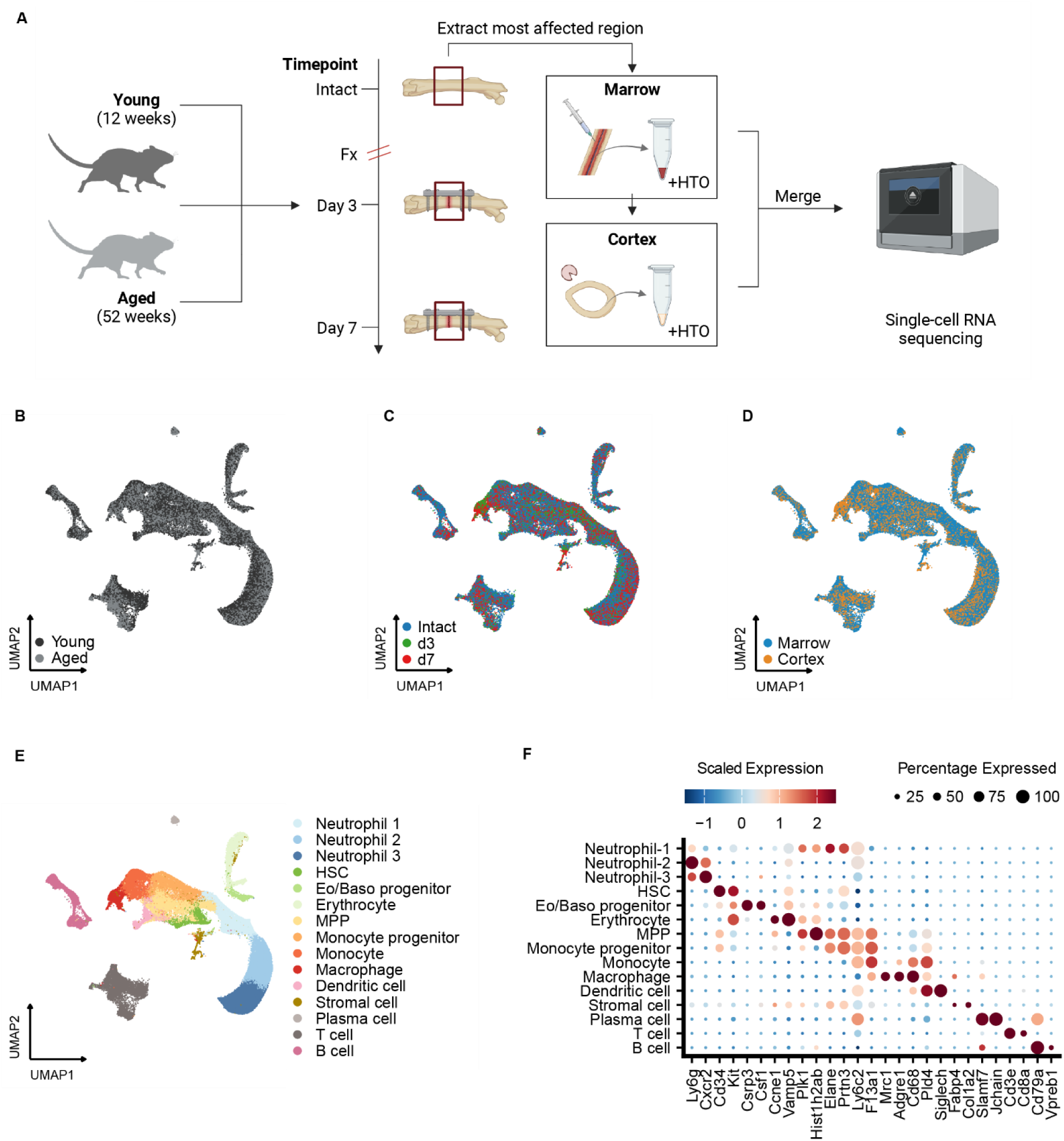
Experimental design and single-cell transcriptomic profiling of the fracture microenvironment. (A) Schematic overview of the *in vivo* fracture model and sample processing workflow. Young (12-week-old) and aged (52-week-old) mice underwent a standardized femoral osteotomy. Fracture callus tissue from the most affected region was harvested at day 3 (d3) and day 7 (d7) post-osteotomy, along with age-matched intact controls. Bone marrow was extracted by flushing the intramedullary canal, while cortical tissue was digested enzymatically. Hashtag oligonucleotide (HTO) labeling was used to retain sample identity prior to pooling and single-cell RNA sequencing. (B-D) UMAP projections of 178,416 single cells derived from marrow and cortex samples across all conditions. Cells are color-coded by age group (B: young and aged), timepoint (C: intact, d3 and d7) and tissue origin (D: marrow and cortex). (E) Cell type annotation of identified clusters based on canonical gene expression markers reveals distinct immune and stromal populations, including innate and adaptive immune cells as well as stromal cells. (F) Dot plot showing the scaled expression of selected marker genes used for cluster annotation. Dot size indicates the percentage of cells expressing each gene per cluster, color intensity reflects average expression level.

In total, we recovered 178,416 cells across all conditions. To identify distinct cell populations among the compartments, we applied uniform manifold approximation and projection (UMAP) analysis for dimensionality reduction and Louvain clustering, which revealed 16 unique clusters (Figure 2B-E). These clusters were annotated based on the expression of well-established marker genes, corresponding to a range of immune and stromal cell types, including 3 neutrophil clusters (Neutrophil-1, Neutrophil-2 and Neutrophil-3), hematopoietic stem cells (HSCs), eosinophil/basophil progenitors, erythrocytes, multipotent progenitors (MPPs), monocyte progenitors, monocytes, macrophages, dendritic cells, stromal cells, plasma cells and T and B cells (Figure 2F). These annotations were validated by cross-referencing established markers from prior single-cell atlases of bone marrow and associated tissues, ensuring consistency with published datasets (Suppl. Figure 1E-F)^25,30–32^.

Following the identification of major cell populations across compartments, we next quantified the relative proportions of each cell type to systematically characterize how fracture injury reshapes the cellular landscape over time. In the marrow, progenitor populations seemed to have one of the most dynamics changes (Suppl. Figure 2). Neutrophil-1, likely resembling neutrophil progenitors (Suppl. Figure 1), and monocyte progenitors expanded significantly at day 3. Notably, these expansions were not sustained as the progenitor levels returned to baseline level at day 7. In addition, mature immune cells also reacted to the osteotomy (Figure 3A). For instance, macrophages showed an increase at day 3 followed by a decline at day 7, while dendritic cells and B cells showed the opposite (Suppl. Figure 2A). These shifts are consistent with an emergency hematopoietic response to recruit innate effectors from the marrow at day 3, and afterwards an inflammatory resolution phase by day 7. At the cortex, the temporal analyses were characterized by an accumulation of innate immune cells (Figure 3B). Macrophages also displayed here an increase at day 3 and a decline at day 7. Furthermore, the Neutrophil-3 cluster expanded over time, reaching their peak at day 7. In contrast to the marrow, the cortex had more cell types, such as dendritic cells and B cells but also plasma cells and erythrocytes, declining in levels during our chosen timepoints (Suppl. Figure 2B). This likely reflected not a loss of these cell types at the cortex, but rather a proportional shift for innate immunity at the cortex after osteotomy.

**Figure 3:**
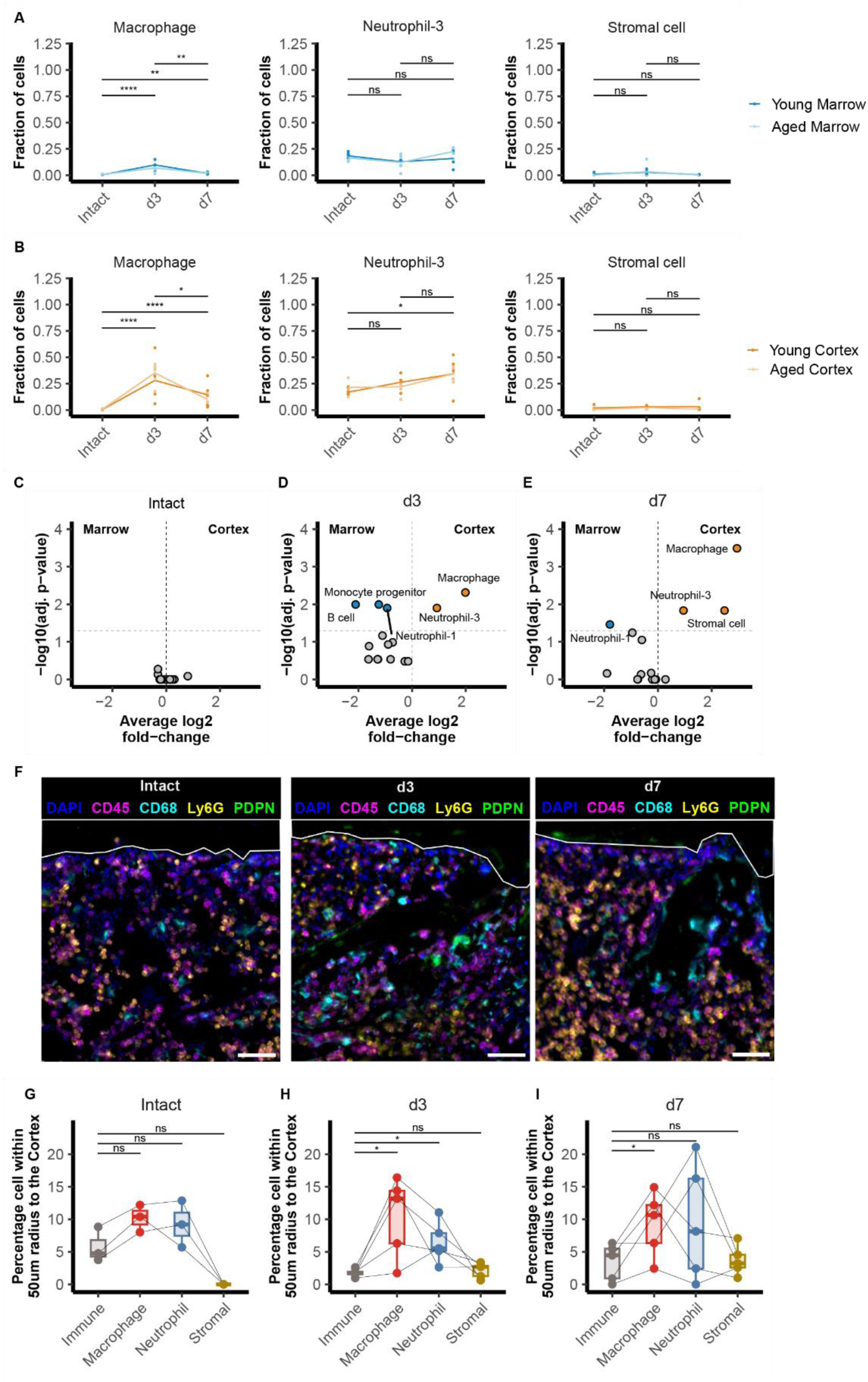
Temporal dynamics and spatial enrichment of macrophages, neutrophils and stromal cells at the cortical bone surface during early fracture healing. (A-B) Proportional abundance of macrophages, neutrophil-3 and stromal cells across timepoints (intact, day 3 and day 7) in marrow (A, blue) and cortex (B, orange). (C-E) Volcano plots illustrating differential enrichment of specific cell populations between marrow and cortex at each timepoint. At day 3 (D), macrophages and neutrophil-3 were significantly enriched at the cortex compared to the marrow. By day 7 (E), the cortical enrichment of macrophages, neutrophil-3 and stromal cells persisted. (F) Immunofluorescence images of intact, day 3 and day 7 samples showing cortical localization of CD45^+^CD68^+^ macrophages (cyan) and CD45^+^LY6G^+^ neutrophils (yellow) in proximity to the cortical bone surface (white line). Nuclei stained with DAPI (blue); CD45 (magenta) marks general immune cells; PDPN (green) indicates stromal cells; The line (white) indicates the cortical surface at the marrow site. Scale bar = 50 µm. (G-I) Quantification of spatial proximity of different cell types to the cortical bone surface, defined as the percentage of each population within a 50 µm radius from the cortex for intact (G), day 3 (H) and day 7 (I). n = 4-5 technical replicates per group; *p < 0.05, **p < 0.01, ****p < 0.0001; ns = not significant, Wilcoxon test with Benjamini-Hochberg correction for multiple testing.

Having characterized the temporal immune and stromal dynamics within each individual compartment, we next compared the distribution of the populations between the marrow and cortex. Cell frequencies were comparable between compartments in intact femurs but changed significantly in a compartment-specific manner post-osteotomy (Figure 3C). At day 3, the marrow displayed an enrichment in progenitors as well as B cells (Figure 3D). In parallel, the cortex exhibited a relative enrichment of Neutrophil-3 cells and macrophages (Figure 3D). By day 7, the distribution of cell populations was further altered between compartments. In the marrow, only Neutrophil-1 remained elevated while monocyte progenitors and B cells were no longer detected (Figure 3E). At the cortex, macrophages and Neutrophil-3 continued to proportionally accumulate compared to their marrow counterparts (Figure 3E). Additionally, stromal cells started to enrich at the cortex, despite limited changes in their overall abundance over time (Figure 3A-B). These results indicated that osteotomy led to a selective recruitment of distinct cell types to the cortex and marrow in both young and aged animals.

Importantly, these injury-associated immune shifts in cell composition were not observed in contralateral bones (Suppl. Figure 3). In contralateral bones, we did not detect a change in frequencies of any immune cell population, including neutrophils, macrophages, and stromal cells, and we detected no substantial difference in the frequency of these cell populations between the marrow and the cortex. These findings collectively highlight that fracture-induced immune responses are highly compartmentalized, governed by localized damage cues rather than systemic effects.

To validate the selective recruitment of macrophages, neutrophils and stromal cells to the cortex following osteotomy, we next employed a 12-marker MELC panel to visualize cell distribution in bone tissue at days 3 and 7 post-osteotomy, with intact femurs from age-matched mice serving as controls (Figure 3F, Suppl. Figure 4). Consistent with our single-cell data, we did not detect an enrichment of either CD68+CD45+ cells (in their majority macrophages) or Ly6G+CD45+ cells (neutrophils) within 50 µm of the cortex in intact bones (Figure 3G). However, at day 3, we observed an enrichment of CD68+CD45+ cells and Ly6G+CD45+ cells compared to other immune cells (Figure 3H). By day 7, the enrichment of CD68+CD45+ cells remained significantly elevated, whereas Ly6G+CD45+ cells only showed a modest abundance near the cortex (Figure 3I). In contrast, PDPN+CD45-cells (stromal cells) were sparsely distributed and remained at low frequency across all conditions (Figure 3G-I), suggesting a more limited involvement in the early fracture response. Together, these data confirm the accumulation of distinct immune cell populations at the cortical interface (but not the marrow) during early fracture healing and support a potential role for spatially organized macrophage and neutrophils activity in the early inflammatory and early osteoclast precursor response.

### Spp1hi macrophages act as cortex-specific precursors in osteoclastogenesis

Given the observed enrichment of macrophages at the cortical niche, where early osteoclastic activity is initiated, we next determined which macrophage subsets act as osteoclast precursors during bone regeneration. Therefore, we further isolated all cells annotated as monocyte progenitors, monocytes, and macrophages from the full scRNAseq dataset and performed a subset-specific dimensionality reduction using UMAP visualization. By using Louvain clustering, we identified 6 distinct subpopulations (Figure 4A-B), which showed a clear separation for cells derived from intact and fractured tissue (Figure 4C).

**Figure 4:**
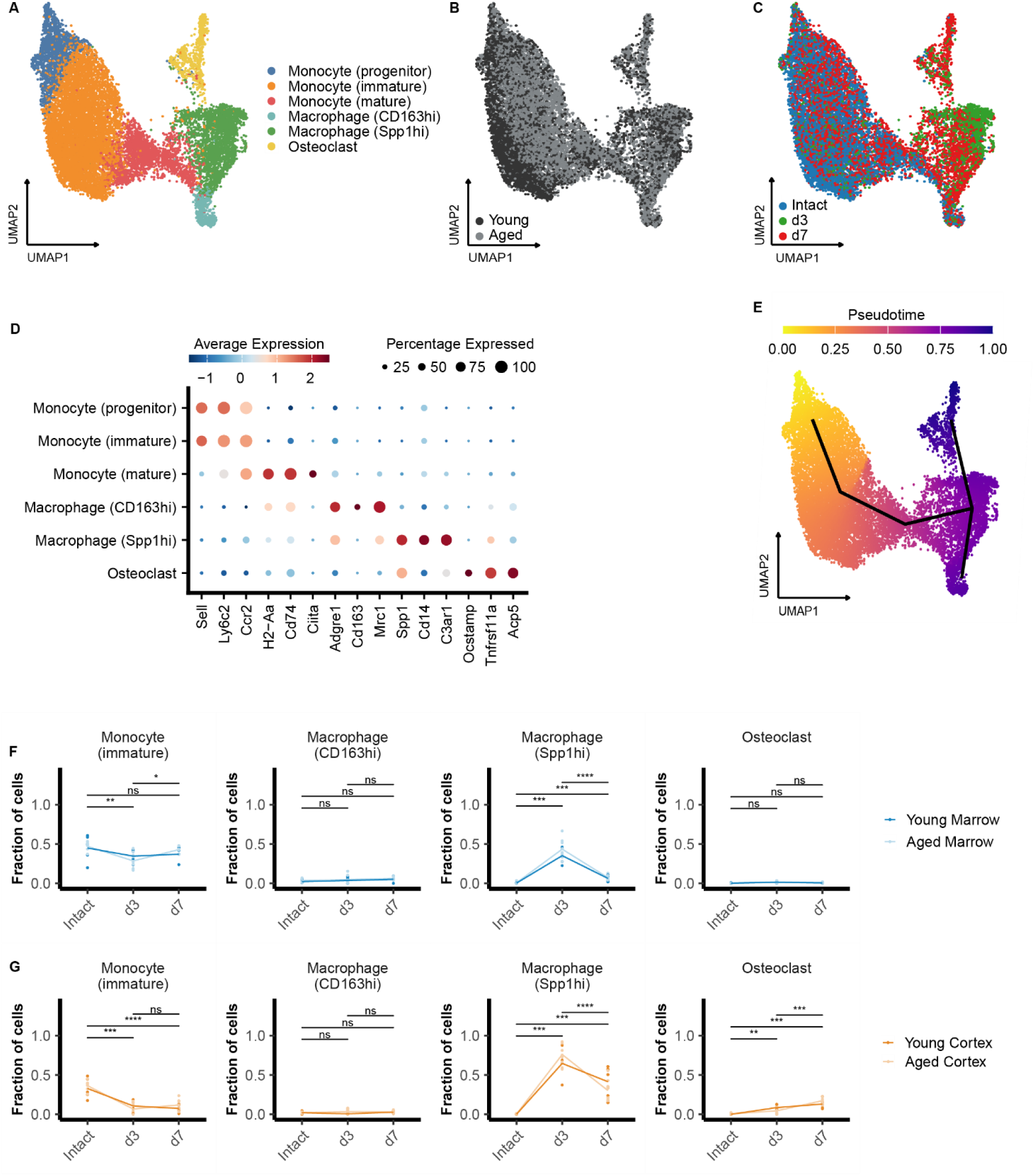
Distinct temporal and tissue-specific differentiation trajectories of monocyte-derived macrophages during early fracture healing. (A) UMAP embedding of macrophage and monocyte lineage cells. (B-C) UMAP projections colored by age group (B: young and aged) and timepoint (C: intact, d3 and d7). (D) Dot plot of representative marker genes used to define each cluster, with dot size representing percentage of expressing cells per cluster and color indicating scaled expression. (E) Pseudotime projection of the macrophage lineage, with two distinct trajectories emerging from monocyte progenitors: one leading toward CD163hi macrophages in the marrow, and another via Spp1hi macrophages toward osteoclast differentiation in the cortex. (F-G) Line plots showing changes in the fraction of each macrophage subtype over time (intact, d3, d7) in marrow (F) and cortex (G). Data points represent individual replicates, with lines connecting group means. n = 4-5 technical replicates per group; *p < 0.05, **p < 0.01, ***p<0.001, ****p < 0.0001; ns = not significant, Wilcoxon test with Benjamini-Hochberg correction for multiple testing.

Based on canonical marker gene expression and cross-referenced with established monocyte-derived macrophage and osteoclast differentiation datasets^24,33,34^, these populations were annotated as monocyte progenitors, immature monocytes, mature monocytes, *Spp1*^hi^ macrophages, *Cd163*^hi^ macrophages as well as osteoclasts (Figure 4D). Monocyte progenitors were defined by expression of *Sell* and high levels of *Ly6c2*, while immature monocytes expressed both *Ly6c2* and *Ccr2*, a chemokine receptor critical for monocyte recruitment to sites of injury. Mature monocytes were characterized by *Ccr2* in combination with *Cd74* and *Ciita*, genes associated with antigen presentation and inflammatory activation. The *Spp1*^hi^ macrophage population showed strong expression of *Adgre1*, *Spp1*, *Cd14*, *C3ar1*, and *Mrc1*. Notably, *Cd14* and *C3ar1* are typically linked to M1-like pro-inflammatory macrophage function^33^, where *Cd14* acts as a co-receptor for TLR signaling^35^ and *C3ar1* is involved in complement-mediated activation^36^. *Spp1*, encoding osteopontin, is associated with fibrosis and tissue remodeling, and while it is often linked to pro-inflammatory processes, its role in fracture healing remains complex^37,38^. On the other hand, *Mrc1,* encoding mannose receptor 1, is a hallmark of M2-like anti-inflammatory and tissue-repairing macrophages. This co-expression suggests that *Spp1*^hi^ macrophages occupy a transitional state bridging inflammatory and reparative phenotypes. *Cd163*^hi^ macrophages were defined by their high expression of *Adgre1*, *Mrc1*, and *Cd163*, and are associated with anti-inflammatory function and tissue remodeling. Finally, osteoclasts expressed canonical bone-resorptive genes *Acp5*, *Tnfrsf11a*, and *Ocstamp*, consistent with their role in matrix degradation and late-stage tissue remodeling.

Having identified osteoclasts among our macrophage subsets, we next performed pseudotime analysis to explore whether monocyte progenitors give rise to osteoclasts through defined intermediate states following osteotomy. The resulting trajectory revealed a continuous progression originating from monocyte progenitors, transitioning through immature and mature monocytes, and branching into either *Cd163*^hi^ macrophages or osteoclasts as terminal fates (Figure 4E). Interestingly, *Spp1*^hi^ macrophages emerged as the key bifurcation point, marking the decision node between a reparative macrophage identity and an osteoclast fate. As osteoclasts did not arise directly from monocytes but only through the *Spp1*^hi^ intermediate, we acknowledge that this population may alternatively be classified as transitional monocytic phagocytes. Nonetheless, these data provide evidence that the transcriptionally distinct population referred to as *Spp1*^hi^ macrophages function as osteoclast precursors *in vivo*.

To investigate whether the trajectories are reflected in the time kinetics of macrophage populations *in vivo*, we next quantified the relative proportions of each subset across timepoints and compartments following fracture (Suppl. Figure 5A-D). In the marrow, both immature monocytes and *Spp1*^hi^ macrophages exhibit dynamic changes (Figure 4F): a drop for the immature monocytes and an increase for *Spp1*^hi^ macrophages at day 3, both returning partially toward baseline by day 7. This pattern suggests a rapid recruitment and differentiation process, likely driven by injury cues. *Cd163*^hi^ macrophages and osteoclasts remained relatively stable over time, underscoring the unique temporal responsiveness of the *Spp1*^hi^ macrophages during the early inflammatory phase.

At the cortex, similar trends but with greater amplitude were observed (Figure 4G). The early surge in *Spp1*^hi^ macrophages and the progressive increase in osteoclast numbers in the same niche point to a localized differentiation process, where *Spp1*^hi^ macrophages accumulate at the injury site and give rise to osteoclasts in situ. The absence of such dynamics in contralateral controls reinforces the conclusion that these events are locally induced and injury-specific (Suppl. Figure 5E).

To further validate this compartmentalized differentiation, we assessed the tissue-specific distribution of these population (Figure 5A, Suppl. Figure 5C-D). After osteotomy, the marrow remained rich in immature and mature monocytes, whereas *Spp1*^hi^ macrophages became enriched specifically in the cortex (Figure 5B). Over time, *Cd163*^hi^ macrophages and osteoclasts were also enriched, but in distinct compartments: *Cd163*^hi^ in the marrow and osteoclasts in the cortex (Figure 5B). This spatial segregation supports the model in which the cortical microenvironment selectively promotes osteoclastogenesis, while the marrow sustains alternative fates.

**Figure 5:**
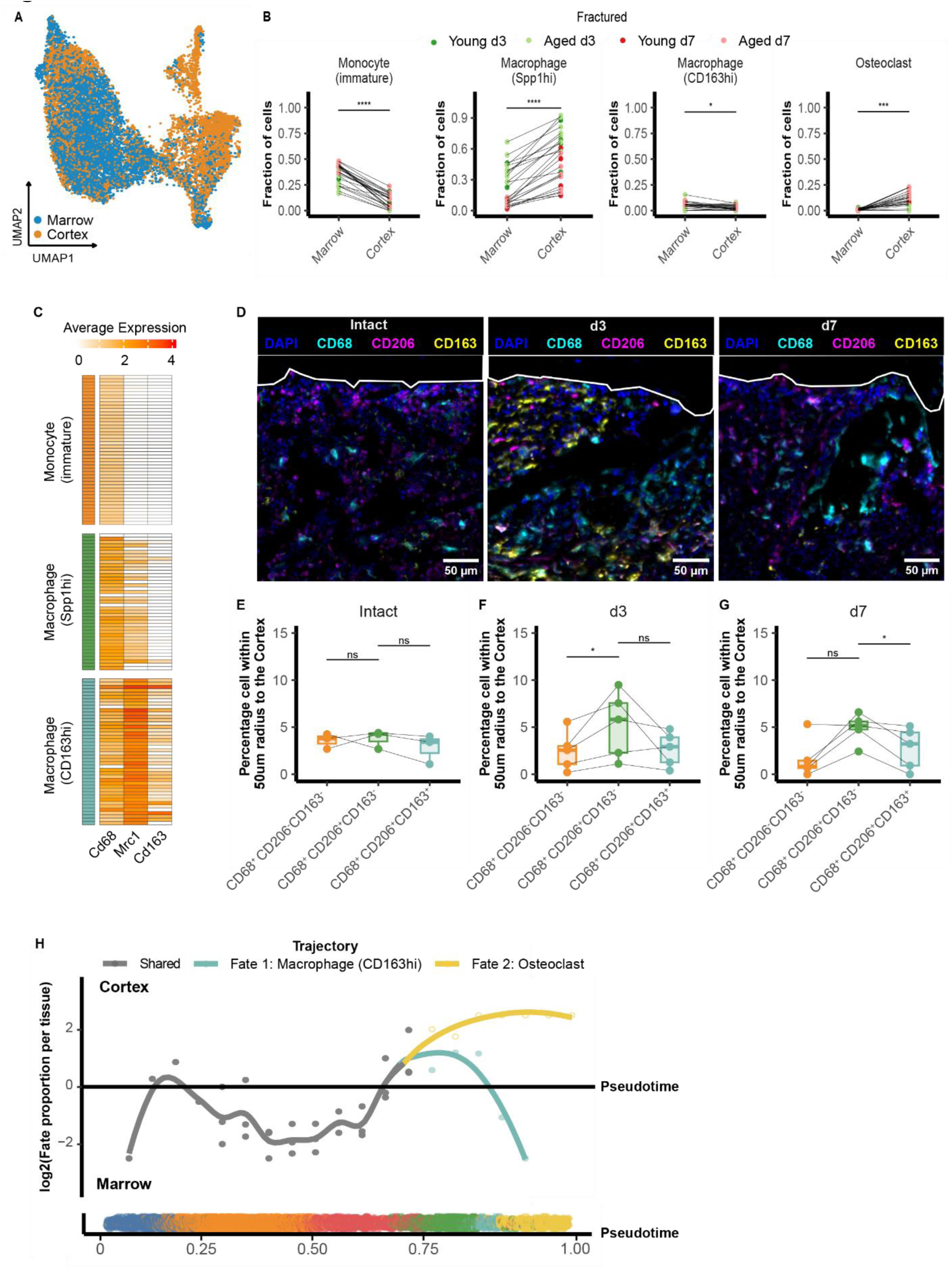
Compartment-specific fate bias of macrophage subtypes during early fracture healing. (A) UMAP embedding of macrophage/monocyte lineage cells from all samples colored by tissue origin (marrow: blue; cortex: orange), revealing spatial segregation of macrophage subtypes. (B) Paired quantification of monocyte and macrophage subtype distributions across marrow and cortex for young and aged mice in fractured conditions (d3 and d7 post-osteotomy). (C) Heatmap showing normalized expression of key markers across macrophage subsets, confirming phenotype-specific gene expression signatures. (D) MELC images showing cortical localization of CD68^+^CD206^-^CD163^-^ monocytes, CD68^+^CD206^+^CD163^-^ macrophages and CD68^+^CD206^+^CD163^+^ macrophages. DAPI (blue) marks nuclei; CD68 (cyan) marks general myeloid cells; CD206 (magenta) and CD163 (yellow) identify functionally distinct subsets. White lines indicates the cortical bone surface; The line (white) indicates the cortical surface at the marrow site. Scale bar = 50 µm. (E-G) Quantification of macrophage subset localization within 50 µm of the cortical surface for intact (E), day 3 (F) and day 7 (G) samples. (H) Pseudotime-aligned plot showing inferred differentiation trajectories of monocyte/macrophage lineage across pseudotime. Log_2_-transformed fate probabilities per tissue are shown along the trajectory. Cells are assigned to one of three inferred fates: shared (gray), CD163hi macrophages (cyan) or osteoclasts (yellow). Colored points below the plot indicates cell-wise trajectory labeling across pseudotime. n = 4-5 technical replicates per group; *p < 0.05, **p < 0.01, ***p<0.001, ****p < 0.0001; ns = not significant, Wilcoxon test with Benjamini-Hochberg correction for multiple testing.

To visualize the enrichment of macrophage subsets, we established a macrophage subset-specific marker panel to distinguish among monocytes, *Spp1*^hi^ macrophages and *Cd163*^hi^ macrophages based on our single-cell data. We found *Cd68, Mrc1* and *Cd163* transcripts to be differentially expressed among the monocyte/macrophage populations (Figure 5C). Using MELC, we confirmed the presence of a CD68^+^CD206^-^CD163^-^ population, consistent with monocytes, a CD68^+^CD206^+^CD163^-^ population, consistent with the Spp1hi macrophage phenotype, and a CD68^+^CD206^+^CD163^+^ population, consistent with the CD163hi macrophage phenotype, in our regions of interest (Figure 5D).

We next assessed the frequency of CD68^+^ monocytes, CD68^+^CD206^+^ macrophages and CD68^+^CD206^+^CD163^+^ macrophages within a radius of 50 µm of the cortex. Consistent with our single-cell results, we observed an enrichment of CD68^+^CD206^+^ macrophages in comparison to the other myeloid populations after injury (Figure 5E-G). CD68^+^CD206^+^CD163^+^ macrophages were also present in proximity to the cortex but constituted a lower percentage of their total population, indicating that while these cells contribute to the early fracture response, they are less dominant in this niche compared to CD68^+^CD206^+^ macrophages. Together, this confirms that *Spp1*^hi^ macrophages not only occupy the osteoclast precursor position transcriptionally, but also CD68^+^CD206^+^ macrophages, consistent with *Spp1*^hi^ macrophages in our single cell data, localize physically at the site of osteoclastogenesis.

To directly link spatial context to fate decisions, we performed a compartment-resolved pseudotime fate bias analysis (Figure 5H). While early progenitors originated broadly, the transition toward osteoclasts occurred specifically within the cortex. Spp1hi macrophages were concentrated at the bifurcation point, particularly within the cortical compartment, and gave rise to either osteoclasts when remaining in the cortex, or CD163hi macrophages when migrating to the marrow. Overall, these findings suggest an association between tissue compartment and fates where the niche might provide directional cues that resolve Spp1hi cells into distinct fates, underscoring the spatial imprinting of osteoclastogenesis.

### Osteoclastogenic programming of Spp1hi macrophages is preserved with age

To explore how this differentiation program is influenced by age, we performed DEG and KEGG pathway analyses in monocyte-derived populations. Immature monocytes showed early expression of inflammation-associated genes at day 3, shifting toward maturation by day 7 (Suppl. Figure 5F). Differential gene expression of Spp1hi macrophages revealed that Spp1hi macrophages at day 3 exhibited a pro-inflammatory and glycolytic gene signature, upregulating genes involved in immune activation and metabolic reprogramming (Figure 6A). By day 7, their expression profile transitioned to oxidative phosphorylation and anti-inflammatory pathways as well as osteoclast differentiation, indicating a functional shift toward tissue remodeling and repair (Figure 6B). These time-dependent changes confirm that Spp1hi macrophages precede and likely enable the emergence of functional osteoclasts during early healing.

**Figure 6:**
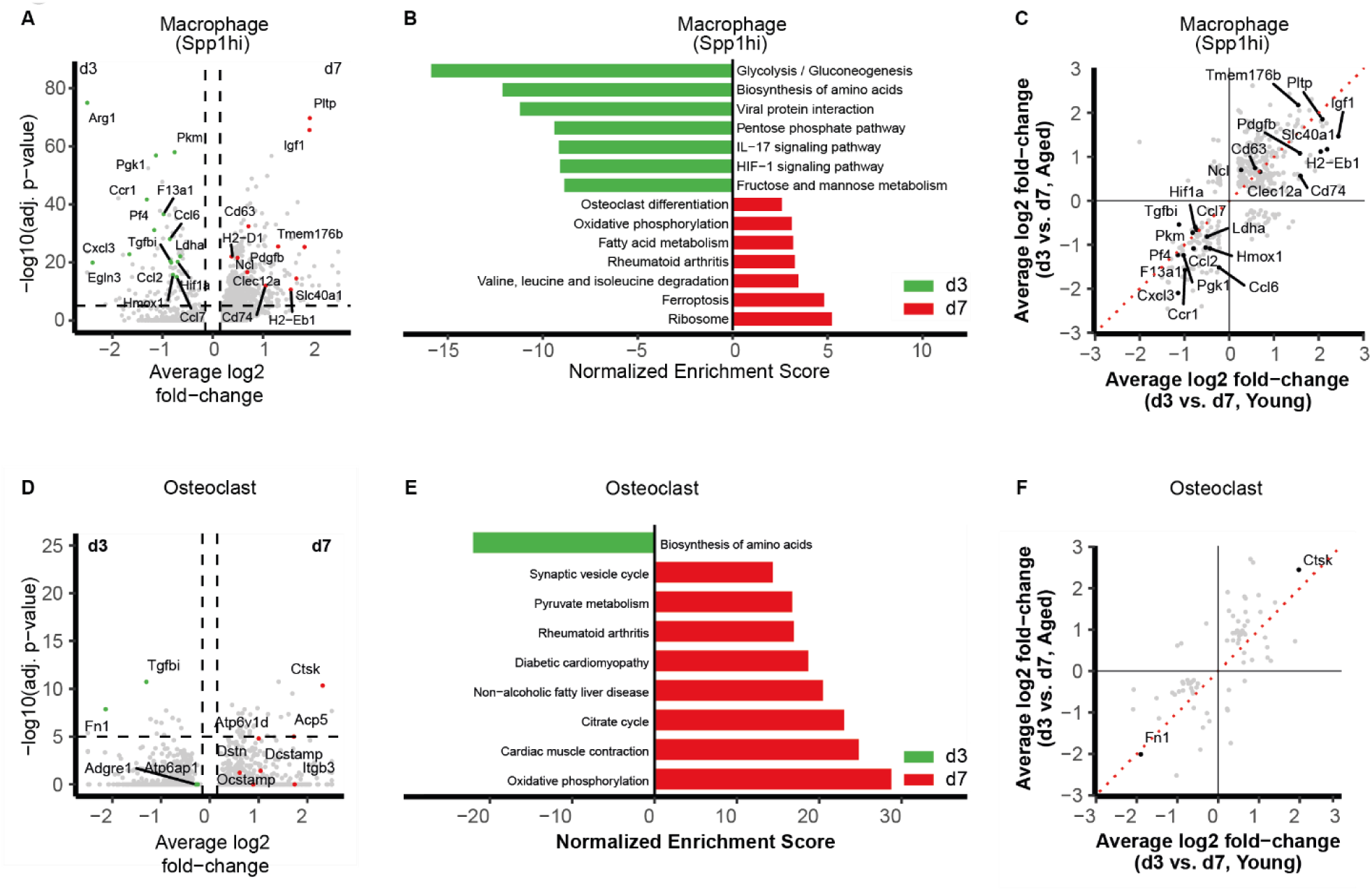
Temporal and age-comparative transcriptomic profiling of Spp1hi macrophages and osteoclasts during fracture healing. (A) Volcano plot showing differentially expressed genes (DEGs) in Spp1hi macrophages between day 3 and day 7 post-osteotomy. Genes labeled in green indicate significant upregulation at day 3; genes in red are upregulated at day 7. (B) KEGG pathway analysis of DEGs in Spp1hi macrophages, comparing day 3 (green) and day 7 (red) based on normalized enrichment scores. (C) Comparative DEG plot for Spp1hi macrophages showing average log^2^ fold change values between day 3 and day 7 in young versus aged mice. Highlighted genes indicate consistent regulation across age. (D) Volcano plot showing DEGs in osteoclasts between day 3 (green) and day 7 (red) post-osteotomy. (E) KEGG pathway analysis of DEGs in osteoclasts comparing day 3 (green) and day 7 (red) based on normalized enrichment scores. (F) Comparative DEG plot for osteoclasts, plotting average log_2_ fold change values for young (x-axis) and aged (y-axis) mice. Black dots represent genes consistently regulated across age.

To further investigate the role of osteoclasts at different stages of fracture healing, we performed differential gene expression analysis to assess their functional state over time (Figure 6D-E). We observed increased expression of *Acp5* and *Ctsk*, both of which are key markers of osteoclast activation and matrix degradation, by day 7 compared to day 3. While osteoclasts were already present at day 3, their transcriptional profile at day 7 suggested an enhanced resorptive function. TRAP staining further supports this transition, as osteoclast activity became more prominent near the cortical fracture site over time (Figure 1), aligning with the observed transcriptional changes. These findings indicate that the cortical microenvironment supports early osteoclast presence but may also provide cues that shape their progressive specialization toward active bone resorption during later stages of healing.

Crucially, these transcriptional programs were largely preserved in aged mice (Figure 6C, F, Suppl. Figure 5G). DEG comparisons revealed that immature monocytes, Spp1hi macrophages, and osteoclasts followed similar activation and differentiation patterns across age groups. While the abundance of these populations may differ, their molecular programs remained intact. This suggests that age-related impairments in osteoclastogenesis are more likely due to altered microenvironmental signals rather than intrinsic cellular dysfunction.

### Neutrophils aid precursor recruitment with subtle age-sensitive niche changes

These findings underscore the central role of the tissue microenvironment in directing immune cell fate during fracture healing. To better understand the nature of these niche signals, we next examined the cellular components that dominate the early immune landscape at the fracture site. In addition to macrophages, Neutrophil-3 cells persisted in the cortical compartment beyond the acute phase (Figure 3B-C), suggesting a more sustained and spatially anchored function. Given the known ability of neutrophils to influence macrophage polarization and osteoimmune signaling via cytokines such as IL-1β, IL-6, IFN-γ, and TNF-α^39^, we hypothesized that neutrophils may contribute to the local regulation of macrophage and osteoclast precursor recruitment and differentiation, including osteoclast formation.

To explore how neutrophils might contribute to niche-specific immune modulation during fracture healing, we investigated their molecular profile over time. Neutrophil-3 exhibited dynamic transcriptional changes over time (Figure 7A-B). At day 3, they expressed pro-inflammatory and immune-activating genes such as *Tnf*, *Csf1*, and *Vegfa*, while by day 7, they upregulated *Mmp8* and genes linked to oxidative phosphorylation and ferroptosis, indicating a functional shift toward matrix remodeling and immune resolution. These temporally regulated states were maintained in aged mice, as gene expression profiles remained highly comparable across age groups (Figure 7C). This suggests that neutrophil maturation and function are preserved with aging, and that their role in shaping the cortical immune niche remains intact.

**Figure 7:**
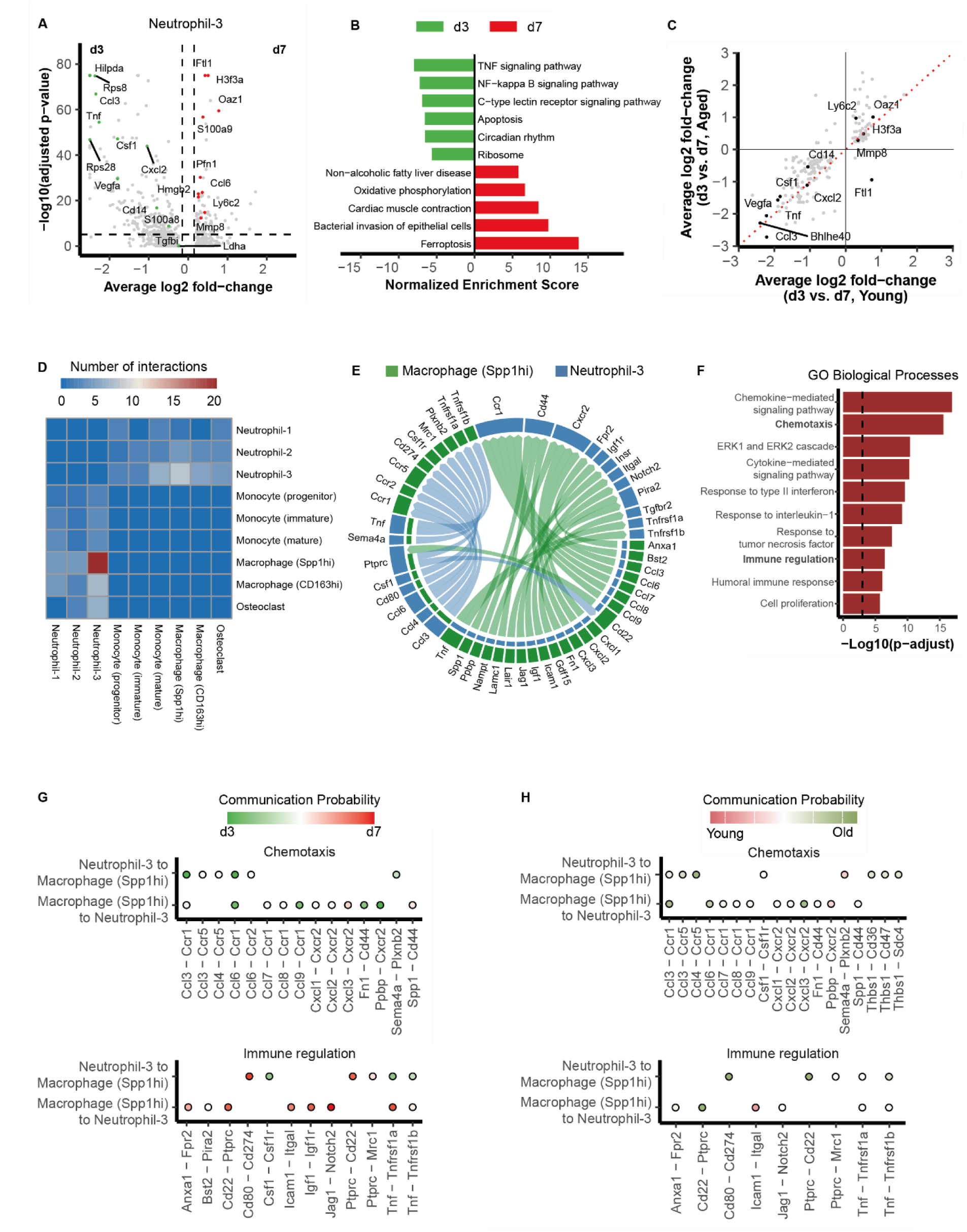
Temporal and directional immune communication between Neutrophil-3 and Spp1hi macrophages during fracture healing. (A) Volcano plot showing differentially expressed genes (DEGs) in Neutrophil-3 cells between day 3 and day 7 post-osteotomy. Genes significantly upregulated at day 3 or day 7 are highlighted in green and red, respectively. (B) KEGG pathways analysis of DEGs from panel A, comparing day 3 (green) and day 7 (red) using normalized enrichment scores. (C) Comparative DEG analysis in Neutrophil-3 cells between young and aged mice, showing average log_2_ fold change for each gene at day 3 vs. day 7. (D) Heatmap displaying the number of identified ligand-receptor pairs between neutrophil and macrophage subtypes across directions and timepoints. Cell color indicates interaction count; columns represent sender cells, whereas rows represent receiver cells. (E) Circos plot visualizing predicted ligand-receptor interactions between Neutrophil-3 and Spp1hi macrophages, stratified by communication direction. (F) Gene ontology (GO) enrichment analysis of ligand-receptor pairs expressed by Neutrophil-3 and Spp1hi macrophages, highlighting processes such as chemotaxis and immune regulation. (G) Dot plots showing temporally resolved ligand-receptor interactions at day 3 and day 7 between Neutrophil-3 and Spp1hi macrophages, grouped by GO biological process: chemotaxis (top row) and immune regulation (bottom row). Dot size reflects interaction score; color represents the dominant timepoint (green: day 3; red: day 7) (H) Dot plots showing ligand-receptor interaction profiles in young and aged mice, across both communication directions and timepoints, grouped by GO biological process: chemotaxis (top row) and immune regulation (bottom row). Dot size reflects interaction score; color represents the dominant age group (red: young; green: aged)

Although neutrophil and macrophage activation programs remained largely intact with aging, we next asked whether their communication within the cortical niche was similarly preserved. Ligand–receptor analysis of cortex-enriched innate immune populations revealed that Neutrophil-3 and *Spp1*^hi^ macrophages formed the dominant cell–cell interaction axis during early fracture repair (Figure 7D). Mapping of ligand–receptor pairs showed a dense signaling network involving key mediators such as *Cxcl2*, *Spp1*, *Tnf*, and *Ccl* family members, linking this axis to both inflammatory recruitment and immune resolution (Figure 7E). GO enrichment further highlighted strong involvement of chemotaxis, cytokine signaling, and regulatory immune pathways (Figure 7F), suggesting that these two populations jointly orchestrate niche dynamics. Together, these findings highlight neutrophil–macrophage communication as a prominent feature of the early spatial immune landscape during bone healing.

We then examined how these interactions evolve over time (Figure 7G). Interestingly, both Cxcl2–Cxcr2 and Spp1–Cd44 signaling remained consistently enriched within the cortical microenvironment across timepoints, suggesting a sustained role in neutrophil recruitment, macrophage activation, and extracellular matrix remodeling^40,41^. However, a temporal shift was apparent in other interactions, with chemotaxis signaling being more dominant at day 3, such as Ccl3-Ccr1, Ccl6-Ccr1 and Ccl9-Ccr1 as well as Fn1-Cd44, while immune regulation pathways gained prominence by day 7, including Cd80-Cd274, Igf1-Igf1r and Jag1-Notch2. Although not directly instructing cell fates, such interactions may reflect mutual priming with neutrophils and macrophages influencing each other’s maturation and potentially guiding macrophages toward osteoclastogenic trajectories. Moreover, this shift suggests that the cortical microenvironment serves as a structured immune niche where neutrophils and macrophages organize their signaling in a time-dependent manner, coordinating the transition from inflammatory recruitment to immune resolution.

Importantly, when comparing young and aged mice, we found that the core signaling framework remained intact (Figure 7H). However, several interactions showed subtle shifts in their relative enrichment. For example, Ccl4-Ccr5 signaling seems to be more present in age cortical compartments, potentially reflecting altered precursor recruitment dynamics^42^. Moreover, Thrombospondin-1 (Thbs1)-associated interactions, which are linked to matrix remodeling^43–45^, were more prominent in aged mice. These differences occurred despite largely preserved transcriptional states in both neutrophils and macrophages, pointing to a subtly rewired cell-cell communication with age.

### Pre-osteoblasts shape an osteoclast-supportive niche at young cortical bone

The cortical microenvironment is, however, not solely shaped by immune cells. Stromal cells, including osteolineage and mesenchymal stromal populations, are well-known sources of growth factors, cytokines, and extracellular matrix components that govern immune cell positioning, survival, and differentiation^46^. To further dissect how the non-hematopoietic compartment contributes to fracture niche regulation and whether this is altered with age, we next focused on the transcriptional dynamics of stromal cells across compartments and timepoints.

Using clustering and canonical marker genes, we identified two broad populations within the stromal compartment: endothelial cells and pre-osteoblasts (Figure 8A-D). While endothelial cells expressed vascular markers such as *Pecam1*, *Emcn*, and *Cdh5*, the pre-osteoblast population was markedly heterogeneous (Figure 8E). Subsets expressing *Vim*, *Col1a1*, *Pdpn*, and *Acta2* likely represent fibroblast- or pericyte-like cells, while others expressed *Acan*, *Sp7*, and *Gpam*, consistent with osteochondral and adipogenic potential. This compositional diversity indicates that multiple stromal lineages coexist in the fracture niche and may functionally support immune coordination and bone repair.

**Figure 8:**
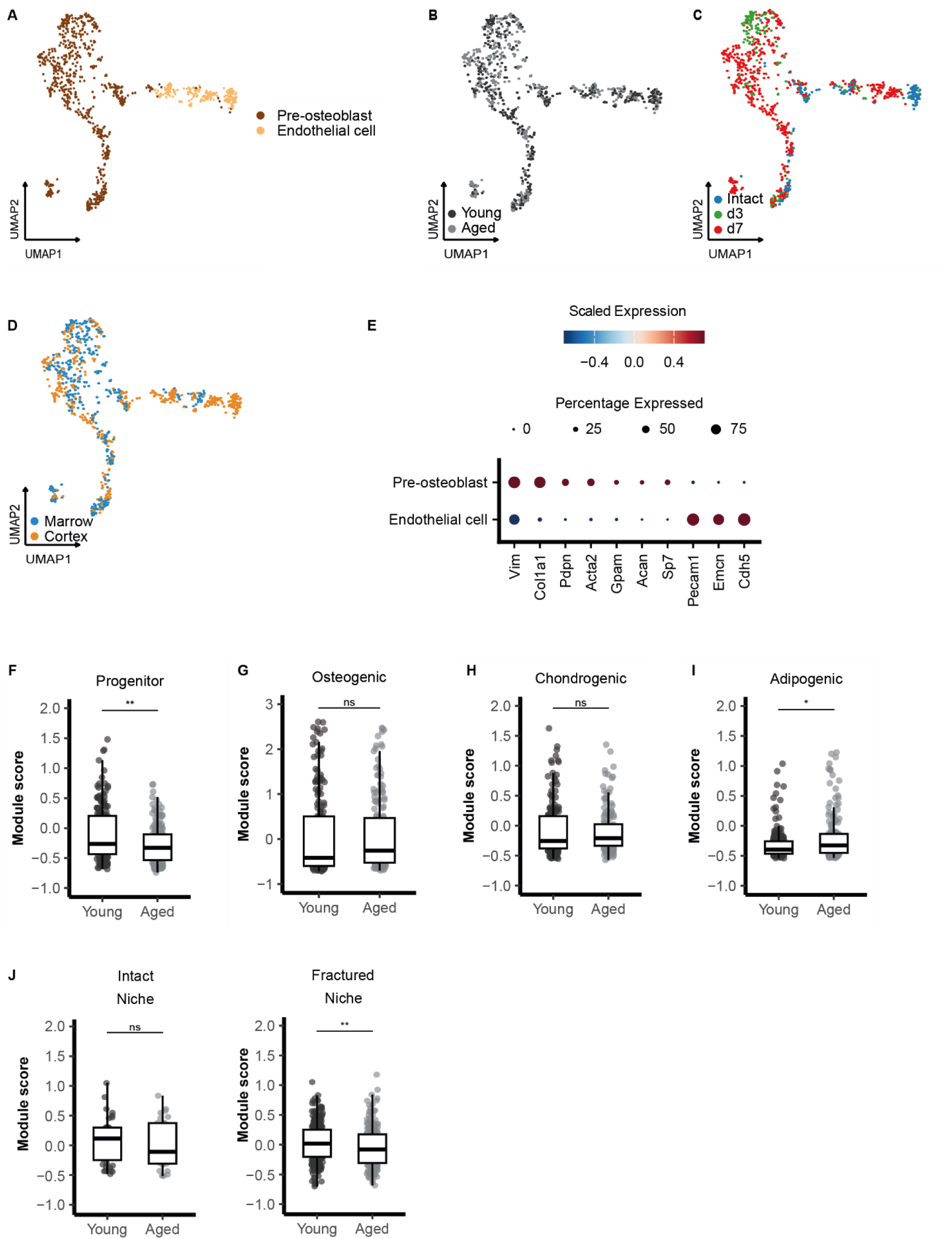
Age-associated changes in progenitor and differentiation potential of pre-osteoblasts. (A-D) UMAP embedding of stromal cells subclustered into pre-osteoblasts and endothelial cells, shown by cell identity (A), age group (B), timepoint (C) and tissue origin (D). (E) Dot plot showing scaled expression and percentage of expressing cells for representative marker genes across the identified stromal subsets, confirming cell type identity. (F-I) Module scores for transcriptional programs related to progenitor identity (F), osteogenic (G), chondrogenic (H) and adipogenic (I) differentiation in young and aged mice. (J-K) Comparison of osteoclastic niche signaling module scores between young and aged mice in intact (J) and fractured (K) tissues. Box plots show median values with interquartile ranges; individual data points represent single cells. *p < 0.05, **p < 0.01; ns = not significant, Wilcoxon test with Benjamini-Hochberg correction for multiple testing.

We next applied gene module scoring to assess the activity of predefined mesenchymal differentiation programs. Pre-osteoblasts from young mice showed higher scores for progenitor-like programs (Figure 8F), indicating a more robust regenerative progenitor pool. Importantly, osteogenic and chondrogenic programs remained relatively comparable between groups (Figure 8G-H), while adipogenic signatures were subtly but consistently lower in young mice (Figure 8I), suggesting a more osteoblast-supportive lineage bias. This balance may enhance the capacity of pre-osteoblast in young mice to contribute to a regenerative fracture environment.

We also assessed whether this regenerative bias was reflected in the expression of niche-regulatory signals essential for immune cell function and osteoclastogenesis (Figure 8J). Pre-osteoblasts from young mice showed higher expression of key factors including *Csf1*, *Tnfsf11* (RANKL), *Cxcl12*, *Wnt5a*, and *Il6*, which collectively mediate myeloid recruitment, survival, and osteoclastogenesis^22^. This elevated niche signaling was most pronounced in the fractured cortical tissue, where young mice maintained higher module scores compared to aged mice. Together, these findings suggest that pre-osteoblasts in young mice provide a more supportive niche environment for osteoclastogenesis during early fracture healing.

## Discussion

Fracture healing is a tightly regulated regenerative process that relies on coordinated interactions between immune and stromal cell populations in a localized and spatially distinct manner. Although osteoclasts are classically associated with late-stage remodeling, evidence points to their very early appearance in bone regeneration and thus an orchestrated recruitment from their precursors directly at the start of healing. However, the cellular origins and regulation of osteoclastogenesis during this early bone repair phase remain incompletely understood. The goal of this study was therefore to identify how tissue microenvironments defines this early process of bone healing, by potentially regulating osteoclastogenesis, eventually altered in aged.

By combining single-cell RNA sequencing with spatial imaging, we reveal that the cortical niche serves as the primary site of innate immune activation, tissue remodeling, macrophage clearing of tissue debris and osteoclastogenesis, while the bone marrow retains its role as a hematopoietic reservoir². This spatial organization of macrophage and osteoclast healing responses were largely preserved in aged animals, indicating that the compartmental framework of fracture repair remains intact with aging. However, despite this preserved structure, we observed an instant increase in early osteoclast number and activity at the fracture site in young mice, distinct from a known increase in baseline osteoclast activity and systemic bone resorption in aged mice^47,48^. This apparent paradox underscores a key distinction between chronic, homeostatic changes with age in bone turnover and the tightly regulated, injury-induced osteoclastogenesis required for regeneration.

To address this discrepancy, we next aimed to identify transcriptional states that may represent osteoclast precursors during early healing. Through transcriptional trajectory analysis, we discovered *Spp1*^hi^ macrophages as a key transitional population, positioned between inflammatory monocytes and terminal osteoclasts. These cells exhibit features of both M1 and M2 macrophages^49,50^; they initially expressing pro-inflammatory markers (M1-like) before transitioning toward a reparative phenotype (M2-like), suggesting they are responsive to local cues that are produced in the cortical microenvironment. Importantly, *Spp1*^hi^ macrophages were specifically enriched in the cortical niche, where osteoclasts also accumulated over time, suggesting in situ differentiation towards osteoclasts. Additionally, their activation trajectory remained largely intact in aged mice, indicating that their intrinsic potential is preserved. Interestingly, similar *Spp1*^hi^ macrophage states have been linked to fibrotic remodeling in other tissues such as the muscle^33^, lung^50^ and kidney^49^. While their role in bone healing appears to favor regeneration, excessive persistence or dysregulation could contribute to pathological outcomes. Together, our findings shed new light to *Spp1*^hi^ macrophage states previously described as fibrosis-related macrophages in other tissues, and now position them as bona fide osteoclast precursors during bone regeneration, whose timely emergence within the cortical niche may influence downstream healing outcomes and offer a potential target for therapeutic intervention.

Our data point to neutrophils as essential catalysts to this macrophage-driven kick-start in the cortical niche at the onset of healing. Traditionally seen as short-lived inflammatory cells, neutrophils persisted in the cortical compartment, transitioning toward matrix-remodeling states. Recent studies have highlighted neutrophil persistence in tissue repair, particularly wound healing and fibrosis models^51–53^, where they influence macrophage activity and ECM remodeling. Similar findings in injury models suggest that dysregulated neutrophil clearance can lead to chronic inflammation^54^, raising the possibility that neutrophil retention in the cortical niche must be tightly regulated to prevent prolonged inflammation. Their predicted ligand–receptor interactions with *Spp1*^hi^ macrophages, particularly via Cxcl2–Cxcr2 and Spp1–Cd44, suggest that also during fracture healing neutrophils and macrophages operate in close coordination, shaping each other’s activation and positioning within the niche. Although neutrophil activation programs were preserved in aged mice, age-specific changes in signaling interactions emerged, such as Ccl4–Ccr5 and Thbs1-mediated pathways. These subtle alterations may reduce the functional relay between neutrophils and macrophages, delaying the osteoclastogenic transition despite preserved cellular states.

While immune-immune interactions coordinate early inflammation and precursor positioning, pre-osteoblasts ultimately provide the niche infrastructure and molecular cues, such as *Csf1*, *Tnfsf11* (RANKL), and *Wnt5a,* required to initiate osteoclast differentiation^55–59^. In our study, these signals were more prominently expressed in young mice, particularly within the fractured cortical compartment, suggesting a more responsive stromal niche early in healing. Gene module analysis further revealed that pre-osteoblasts in young mice exhibited stronger progenitor-like programs and lower adipogenic expression, consistent with a regenerative lineage bias^7,60^. These findings suggest that pre-osteoblasts in young mice actively provide the niche cues necessary to support osteoclast precursor maturation, including signals that coordinate with recruited macrophages and neutrophils. This early stromal activity appears reduced in aged tissue, where the same cues are less effectively engaged following injury. In this context, the ability of the young stromal niche to initiate early remodeling emerges as a second key determinant of regenerative success. Therapeutically enhancing or mimicking these stromal signals may offer a targeted strategy to promote osteoclast function during fracture healing in aging bone.

Several other factors may influence the early cellular events that enable the macrophage to osteoclast activation. In young mice, we observed a clear increase in osteoclast activity at day 7 post-osteotomy, suggesting that local and systemic conditions at this early stage support timeline osteoclastogenesis. In contrast, even subtle shifts in the aged microenvironment may compromise this early response. For example, alterations in vascular or matrix composition may indirectly affect oxygen and nutrient diffusion, thereby influencing the metabolic activation of osteoclast precursors^61,62,63,64^. In addition, B cells, well-established sources of RANKL, were not addressed in this study, but their known role in promoting osteoclastogenesis suggests that alterations in B cell abundance or function with age may also impact the regenerative capacity of the fracture niche^65,66^. Taken together, these factors may not block osteoclastogenesis, but could delay its onset in aged individuals, shifting the regenerative trajectory from the very beginning of the healing process. Future studies should aim to integrate these systemic and cellular components to build a more comprehensive understanding of how aging impairs bone regeneration through altered osteoclastogenesis.

In conclusion, we demonstrate that early osteoclastogenesis from distinct sets of macrophages is not solely a function of precursor identity, but is critically governed by spatially organized niches and their signals involving specific macrophage subsets, neutrophils, and stromal cells. Aging does not abrogate the potential for osteoclast differentiation but alters or disrupts the niche required to support it. Notably, we identify Spp1hi macrophages as early, cortex-restricted transitional cells that defy classical M1/M2 polarization and emerge in close coordination with neutrophils. This co-localized immune module forms a pre-regenerative microenvironment that initiates remodeling cascades from the earliest stages of healing. Its spatial restriction to the cortical compartment, and its absence in intact tissue, underscores its specificity and potential as a checkpoint for successful regeneration. These findings provide a new conceptual framework for how tissue context shapes immune function at the very early start of regeneration, and highlights differences in aged that may be conceptually addressed to improve healing across aging.

## Material and methods

### Animal Model

Female C57BL/6NRj mice were purchased from Janvier with an age of 10 and 26-32 weeks, and were used at an age of 12 and 52 weeks. Animals were imported with a health certificate and kept under obligatory hygiene standards that were monitored according to the FELASA standards. The mice were kept under specific pathogen free (SPF) housing. Food and water was available *ad libitum* and the temperature controlled with a 12h light/dark cycle. All experiments were carried out with ethical permission according to the policies and principles established by the Animal Welfare Act, the National Institutes of Health Guide for Care and Use of Laboratory Animals, and the National Animal Welfare Guidelines, the ARRIVE guidelines and were approved by the local legal representative animal rights protection authorities (G0012/22; Landesamt für Gesundheit und Soziales Berlin).

### Mouse Osteotomy as a Model to Study Bone Regeneration

Bone regeneration was studied with a mouse osteotomy model. The osteotomy was performed under isoflurane anesthesia. The anesthesia was given to the mouse through a mask with an isoflurane-oxygen mixture. Before the surgery, mice were subcutaneously injected with Buprenorphine (analgesic) and Clindamycin (antibiotics), and eye ointment was applied to protect the eyes. The surgical area was shaved and the surgery was performed on a heating pad (37°C). The osteotomy was performed as previously published. To summarize, after shaving the surgical area of the left femur, the skin was opened, and the femur was carefully exposed. An external rigid fixator (MouseExFix, RISystem, Davos, Switzerland) was mounted on the femur and the osteotomy was performed with a gap size of 0.7 mm. Afterwards, the wound was closed and Tramadol was added to the drinking water for 3 days. Animals were sacrificed 3, 7, and 21 days post-surgery through intraperitoneally injection with a mixture of Medetomidine and Ketamine to induce a deep anesthesia, followed by a cervical dislocation.

### Bone Tissue Sample Preparation for Single-cell RNA Sequencing

The femurs were carefully excised and stored in ice cold phosphate-buffered saline (PBS) supplemented with 2% fetal calf serum (FCS) to preserve cellular viability during transportation. For bone marrow isolation, the femurs were sectioned from inner pin to inner pin, followed by flushing the marrow out of the cavity. The resulting cell suspension was filtered through a cell strainer and the red blood cells were lysed with ACK lysis (ThermoFischer Scientific). The reaction was neutralized by adding PBS+FCS solution. To isolate cells from the cortical bone, the bone cavities were washed three times with 10 mL digestion medium (1 mg/mL of each collagenase IV and dispase II in Hanks’ balanced salt solution; all from Gibco) for 30 min at 37°C in a water bath. Following incubation, the digest was filtered through a cell strainer to obtain a cell suspension suitable for further analysis.

### Single-cell RNA Sequencing – 10x Genomics

For scRNAseq using 10x Genomics, marrow and cortical bone cells were processed as described earlier. Each compartment was stained with Cell Hashing antibodies following the CITE-seq protocol. After staining and washing, a final concentration of ∼1000 cells/uL, was loaded into the 10x Genomics Chromium Controller for Single Cell 3’ v3 workflow, according to the manufacturer’s instructions.

In brief, reverse transcription and library construction were carried out per the 10x Genomics protocol. Total cDNA synthesis involved 12-14 amplification cycles, resulting in final cDNA yields ranging from approximately 1 ng/uL and 64 ng/uL. The 10x Genomics sequencing libraries were constructed as described and sequenced on an Illumina NextSeq 500. Sequencing was performed with paired-end read lengths of 26 bp for Read 1 and 58 bp or 98 bp for Read 2.

The raw sequencing reads were processed using Cellranger count (v7.1.0) (10X Genomics), which involved alignment to the mouse genome (mm10) and the generation of count matrices containing both mRNA and HTO read counts. The resulting count data was then analyzed following standard processing pipelines in Seurat^67^ (v.5.1.0). Cells with fewer than 500 detected genes, more than 10% mitochondrial gene expression or identified multiplets were excluded. Cell cycle scores were computed using the CellCycleScoring function, and data were normalized and scaled using NormalizeData and ScaleData functions, which regresses out mitochondrial content, cell cycle phase score and RNA feature counts. Data integration across compartments was performed using Harmony^68^. Dimensionality reduction was carried out using principal component analysis (PCA), with the number of PCs determined by elbow plot inspection. Two-dimensional embeddings were generated using UMAP. Clusters were identified via Louvain clustering (FIndClusters), and differential gene expression was assessed using the FindMarkers function. Iterative subclustering was applied to refine immune populations, and clusters with gene signatures consistent with low-quality cells or contaminating populations were removed.

### Bone Sample Preparation for (immuno)histological analysis

Osteotomized femurs were cryo-embedded according to the Kawamoto method. The harvested bones were directly fixed in 4% paraformaldehyde for 2 h at 4 ^O^C. They were then transferred to a series of increasing sucrose solutions (10%, 20% and 30%) for cryoprotection, with each step lasting 24 hours at 4 ^O^C. After cryoprotection, bones were embedded in SCEM media (SectionLab, Yokohama, Japan) using precooled n-hexane. Embedded bones were stored at −80 ^O^C until sectioning.

For sectioning, the cryo-embedded blocks were mounted onto a cryotome sample holder using Tissue-Tek O.C.T. Compound (Sakura, #4583). Sections of 7 um thickness were cut using a Leica cryostat (Leica), and transferred onto Kawamoto tape (Section lab, #C-FP096). These sections were mounted on a glass slide and fixed with Tesafilm strips for further processing.

### Histomorphometry analysis

For histomorphometry analysis, Movat Pentachrome staining was performed on bone sections. The sections were stained sequentially with alcian blue (8GS, Chroma), Weigert’s haematoxylin (Merck), Brilliant Crocein/Acid Fuchsin (Chroma/Merck), 5% phosphotungstic acid (Chroma), and Saffron du Gâtinais (Chroma). After staining, the sections were mounted using Vitro-Clud (Langenbrinck).

For TRAP staining, sections were fixed in 4% PFA, adjusted to pH 5 using sodium acetate and sodium tartrate, and stained using Naphtol AS-MIX-Phosphate and Fast Red Violet LB Salt. A counterstain was carried out with Mayer’s Hemalaun solution (Merck). TRAP-positive cells with ≥ 3 nuclei, adherent to the bone surface, were identified as osteoclasts.

Mosaic images were captured using an Leica DM6B microscope with Leica LASX Office software (v.1.4.7). Static and cellular histomorphometry parameters were analyzed using ImageJ, following the guidelines of the American Society for Bone and Mineral Research.

### Tissue preparation for MELC

Samples were prepared as described for histologic analysis. Regions of interest were chosen using consecutive slides of each sample stained with Movat Pentachrome. Cuts were thawed on the day of the MELC experiment. The previously identified region of interest of up to 6 samples were cut and glued on a cover slide (24 × 60 mm; Menzel-Gläser, Braunschweig, Germany). The fluid chamber holding 100 μl of PBS was created using “press-to-seal” silicone sheets (Life technologies, Carlsbad, California, USA; 1.0 mm thickness). Permeabilization was performed with 0.2% Triton X-100 in PBS for 10 min at room temperature.

### MELC image acquisition

MELC data acquisition of the osteotomy and the healthy control samples was done using either BioDecipher® Device 1.0 (BioDecipher GmbH, Magdeburg, Germany) or a modified Toponome Image Cycler® MM3 (TIC, (originally produced by MelTec GmbH & Co.KG, Magdeburg, Germany)^69^. The robot and microscope were controlled using the MelTec TIC-Control software.

Antibodies were titrated in order to find the best working concentration in murine bone marrow tissue with detailed information provided in Supplementary Table S1 which also contains the staining order and used acquisition parameters for MELC. Prior to each MELC experiment, PBS with 5 % MACS BSA (Miltenyi Biotec, Bergisch Gladbach, Germany) was prepared and used as washing buffer, and Köhler illumination was performed.

### Image preprocessing

Images were preprocessed using the TIC OBSERVER software (BioDecipher GmbH, Magdeburg, Germany) and as described in Pascual-Reguant *et al.* (2021)^70^. Image normalization was performed with ImageJ^71^ as previously described in Pascual-Reguant *et al*. (2021)^70^.

### Cell segmentation and annotation

Image analysis and cell segmentation were performed using CellProfiler (v.4.2.8). Image intensities were normalized to a scale ranging from 0 to 1. Nuclei were initially identified based on 4’,6-diamidino-2-phenylindole (DAPI) staining, with an expansion step applied to approximate cell size more accurately. The median fluorescence intensity (MFI) of all markers was quantified within each individual cell and exported as a CSV file for further analysis (Supplementary Table S2).

Extracted features were transformed using a hyperbolic arcsine transformation (cofactor = 0.2). Batch correction was implemented using Harmony algorithm allowing data integration across different animals and timepoints. Cell clustering and annotation were performed using DittoSeq^72^. Marker expression was visualized using UMAP, and annotated clusters were validated by overlaying IF images. Spatial proximity analysis was carried out using SPIAT^73^ package, applying average_percentage_of_cells_within_radius function to assess spatial proximity of cells to the cortex.

### Statistical analysis

All statistical analysis were performed using R software (v.4.3.3) The choice of parametric and nonparametric tests is indicated in the figure legends, depending on the data distribution and experimental design. P-values of less than 0.05 were considered statistically significant. For differential gene expression analysis, genes with adjusted p-values of less than 0.001 and an average log-fold change greater than 0.25 were considered significantly expressed. Multiple testing correction was performed using the Benjamini-Hochberg method to control for false discovery rates.

## Data availability

Single-cell RNA-seq datasets (raw FASTQ files and processed count matrices) generated in this study are available at the Gene Expression Omnibus under accession number GSE305266. All Seurat objects as well as all MELC data are available in the Zenodo open-access repository (https://zenodo.org/): the Seurat objects are deposited under https://doi.org/10.5281/zenodo.16871641, the MELC images of intact controls can be found under https://doi.org/10.5281/zenodo.16870991, the MELC image of young animals at day 3 under https://doi.org/10.5281/zenodo.16870196, and the MELC images of young animals at day 7 under https://doi.org/10.5281/zenodo.16871121. Finally, the CellProfiler pipeline and its respective output files can be found under https://doi.org/10.5281/zenodo.16876039. All Zenodo records are interlinked within the public repository and can be accessed directly via their respective DOIs.

## Acknowledgements

We are grateful for the excellent technical assistance of Gabriela Korus, Mario Thiele and Katrin Vogt. We also thank the BIH/MDC Genomics Technology Platform and the Core Unit Bioinformatics for their support and expertise in single-cell sequencing and data analysis.

## Contributions

AN has participated in conception and design of the study, acquisition and analysis of all data and manuscript writing. HZO contributed to single-cell study design and data acquisition. DMM participated in analysis of all data, and manuscript writing. SK contributed to the design of the MELC experiments and was involved in MELC data acquisition and analysis. AE, MK and KSB contributed to the design and coordination of the animal studies. OB has participated in analysis of single-cell data. RU and RG conducted the MELC experiments. CHB, AT, SH and KSB provided conceptual input and contributed to discussions throughout the project. AEH, BS, GND contributed to conception and design of the study. All authors contributed to data interpretation and manuscript editing, and approved the final version of the manuscript.

## Conflict of interest

The authors declare no conflict of interest.

## Funding

This work was supported by funding within the German Research Foundation (DFG), CRC 1444, P13 to BS and GND, and P14 to AEH, as well as ERC Advanced Grant “Immuno-Mechanics”.

## Notes

### Competing Interest Statement

The authors have declared no competing interest.

