## Supplemental Figures for "The cortical microenvironment drives early immune organization and controls early osteoclastogenesis in bone healing"

*Correspondence:

### Supplementary figures

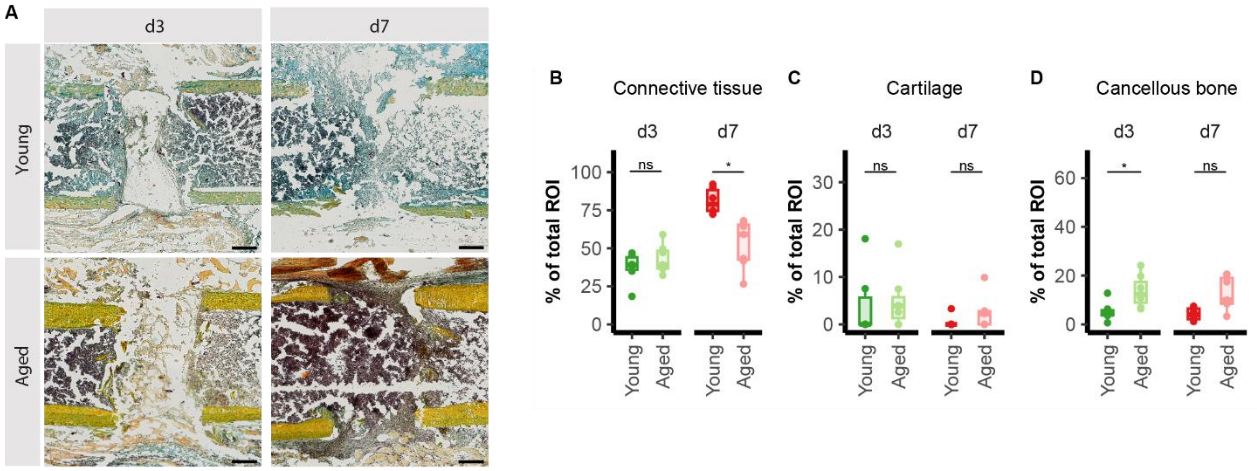


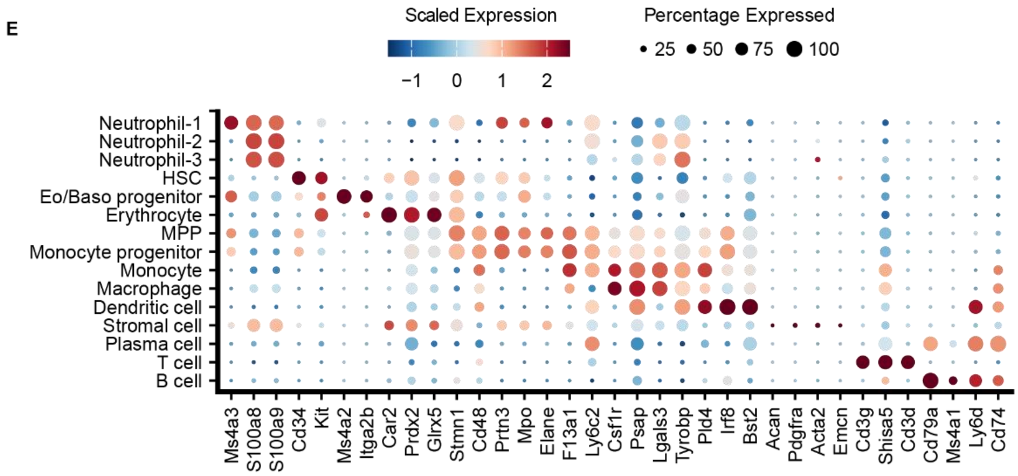


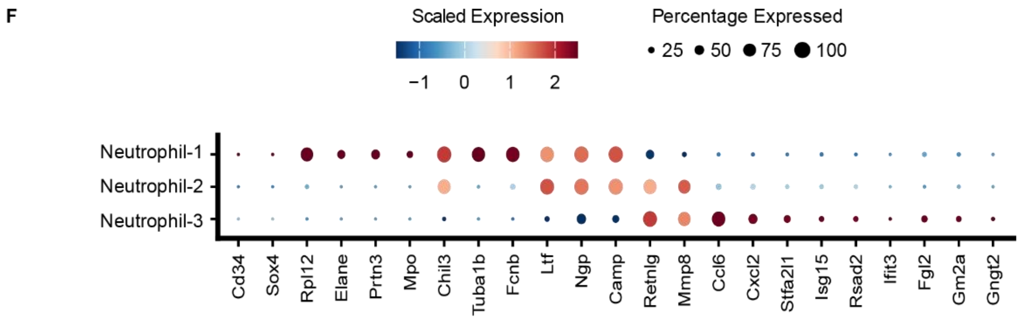

**Suppl. Fig 1: Validation of the *in vivo* osteotomy model for early-phase fracture healing.**

(A) Representative histological sections stained with Movat Pentachrome from young and aged mice at day 3 and day 7 post-osteotomy. Tissue components are visualized as follows: yellow = mineralized bone, red = connective tissue, and blue/green = cartilage. Sections are shown in standardized orientation (proximal to the left, distal to the right). Scale bar = 200 μm.

(B-D) Quantification of the relative area occupied by connective tissue (B), cartilage (C) and cancellous bone (D) within the defined region of interest (ROI = 0.7 mm fracture gap including 0.4 mm proximal and distal). Tissue segmentation was performed using a custom Fiji-based pipeline.

(E-F) Dot plots displaying scaled expression and percentage of expressing cells for gene markers obtained from literature.

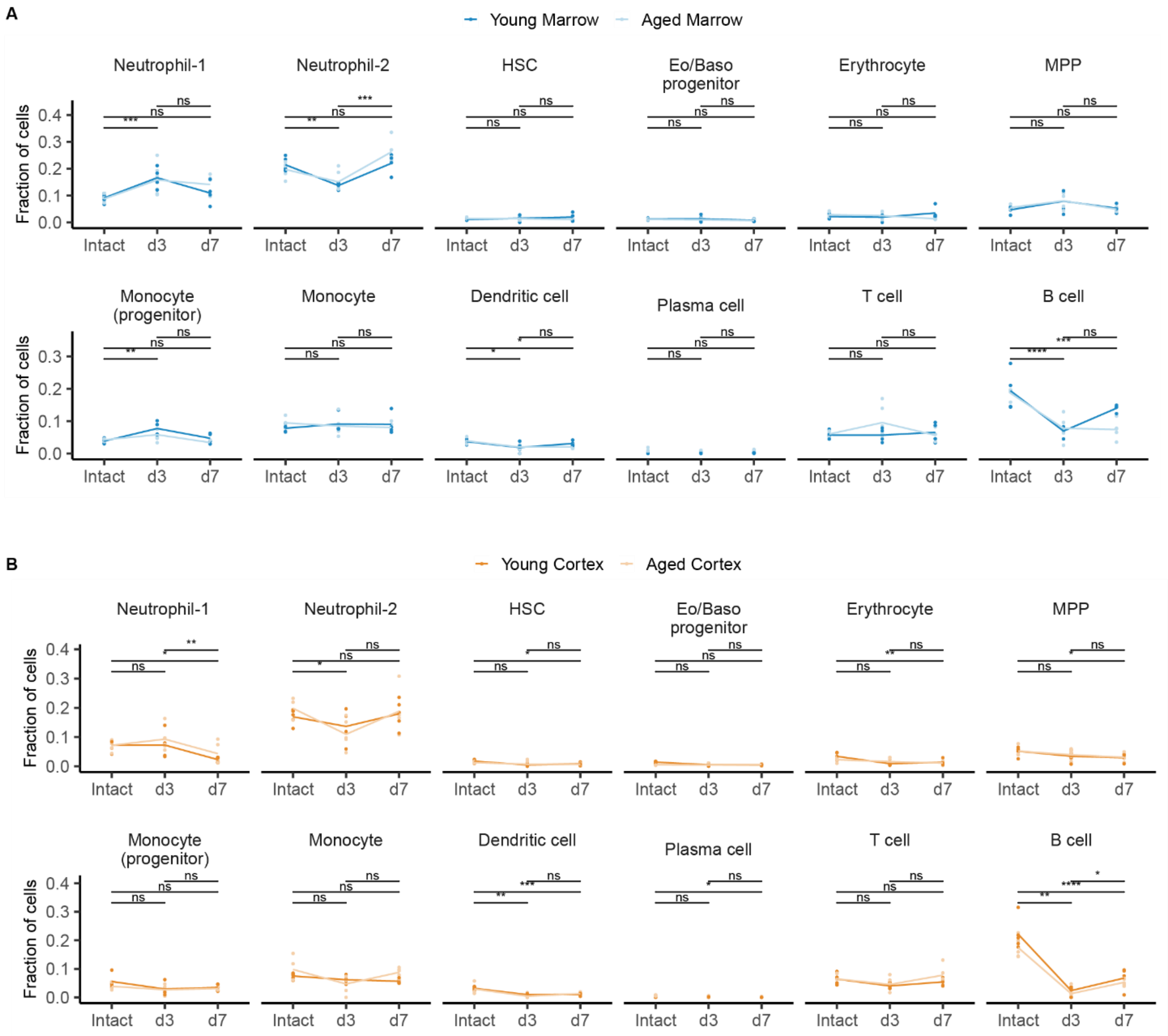


**Suppl. Fig. 2: Temporal distribution of hematopoietic and immune populations during early fracture healing in marrow and cortex.**

Fraction of cells per annotated population at baseline (intact) and post-osteotomy (day 3 and day 7) in marrow (A, blue) and cortex (B, orange). Cell types include Neutrophil-1, Neutrophil-2, hematopoietic stem cells (HSCs), eosinophil/basophil (Eo/Baso) progenitors, erythrocytes, multipotent progenitors (MPPs), monocyte progenitors, monocytes, dendritic cells, plasma cells and T and B cells. Statistical comparisons were performed using Wilcoxon test with Benjamini-Hochberg correction. Each data point represents an technical replicate (n = 4-5 per group). *$p<0.05$, **$p <0.01$, ***$p < 0.001$; ns = not significant.

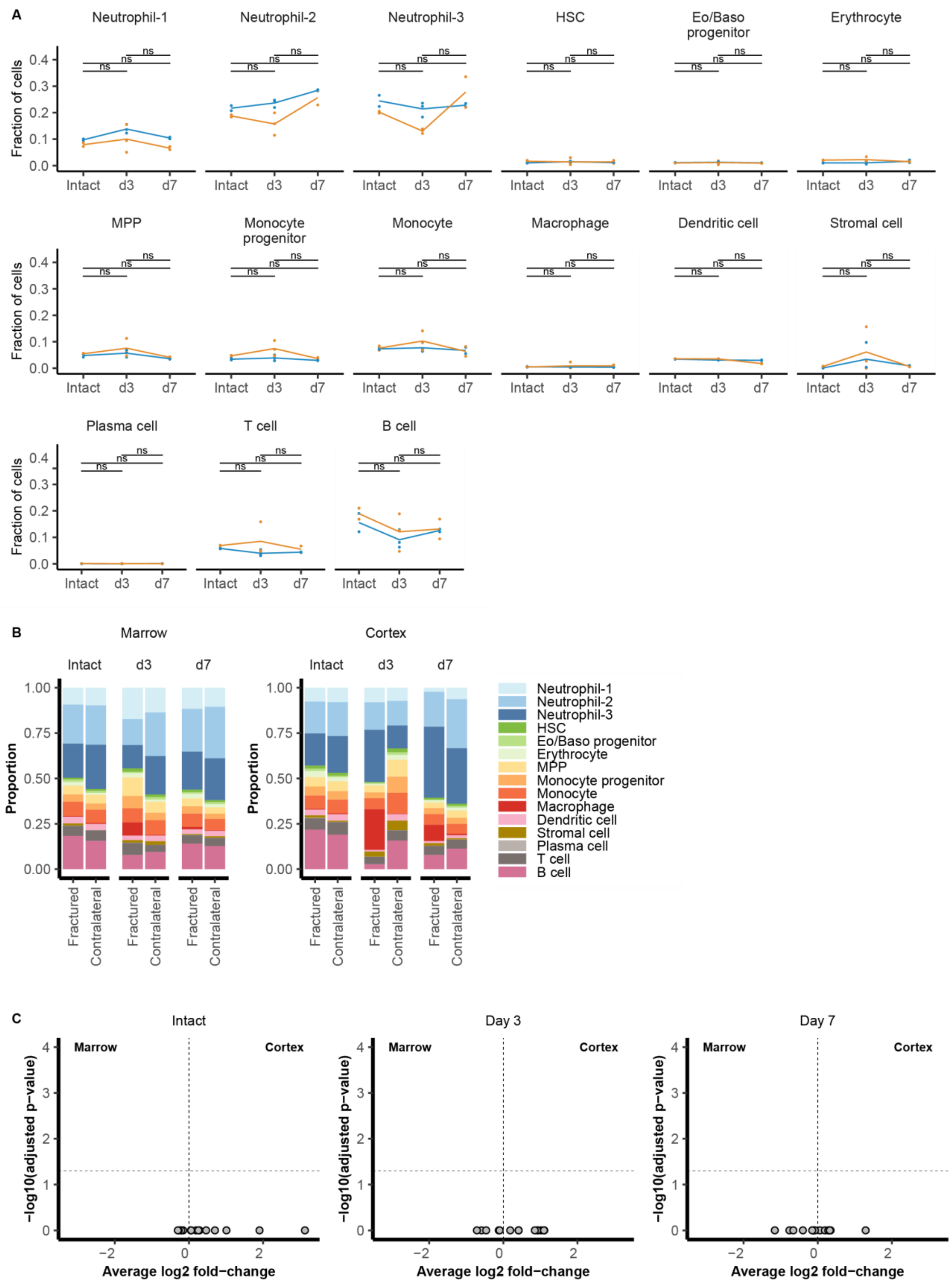
A
Neutrophil-1
Neutrophil-2
Neutrophil-3
HSC
Eo/Baso progenitor
Erythrocyte
MPP
Monocyte progenitor
Monocyte
Macrophage
Dendritic cell
Stromal cell
Plasma cell
T cell
B cell
Fraction of cells
0.0
0.1
0.2
0.3
0.4
ns
Intact
d3
d7
B
Marrow
Cortex
Intact
d3
d7
Proportion
0.00
0.25
0.50
0.75
1.00
Fractured
Contralateral
Neutrophil-1
Neutrophil-2
Neutrophil-3
HSC
Eo/Baso progenitor
Erythrocyte
MPP
Monocyte progenitor
Monocyte
Macrophage
Dendritic cell
Stromal cell
Plasma cell
T cell
B cell
C
Intact
Day 3
Day 7
Marrow
Cortex
−log10(adjusted p−value)
0
1
2
3
4
−2
0
2
Average log2 fold−change

**Suppl. Fig. 3: Stable baseline composition and compartmental consistency across timepoints and tissue locations.**

(A) Proportional abundance of annotated populations across timepoints (intact, day 3 and day 7) in marrow (A, blue) and cortex (B, orange) of contralateral bones.

(B) Stacked bar plots showing proportional distribution of annotated cell types in fractured and contralateral samples from marrow and cortex compartments at each timepoint.

(C) Volcano plots showing differential abundance analysis of cell types between fractured and contralateral sides in marrow and cortex. No significant differences were detected across timepoints in the absence of injury, supporting the use of contralateral tissue as an internal control.

n = 2-3 technical replicates per group; *$p < 0.05$, **$p < 0.01$, ****$p < 0.0001$; ns = not significant, Wilcoxon test with Benjamini-Hochberg correction for multiple testing.

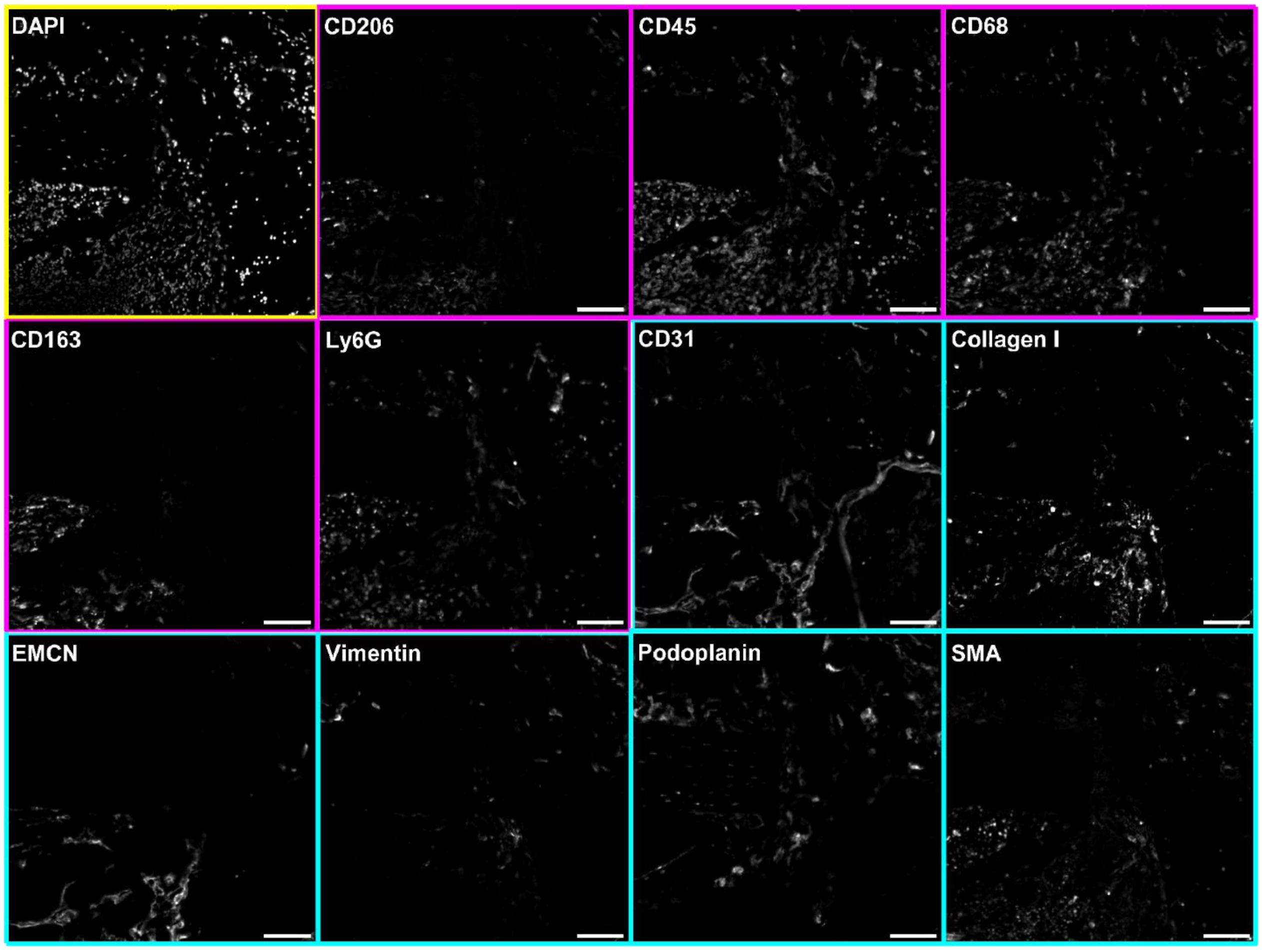


**Suppl. Fig. 4: MELC panel for cortical bone tissue**
Representative field of view from bone tissue near the cortical surface, sequentially stained using a 12-marker MELC panel. Markers include immune cell populations (magenta), stromal and endothelial cell markers (cyan) and nuclei stainings (yellow). Each marker was visualized in the same tissue region through cyclic immunofluorescence. Image resolution is 2048x2048 pixels, corresponding to 665x665 µm. Scale bar: 100 µm.

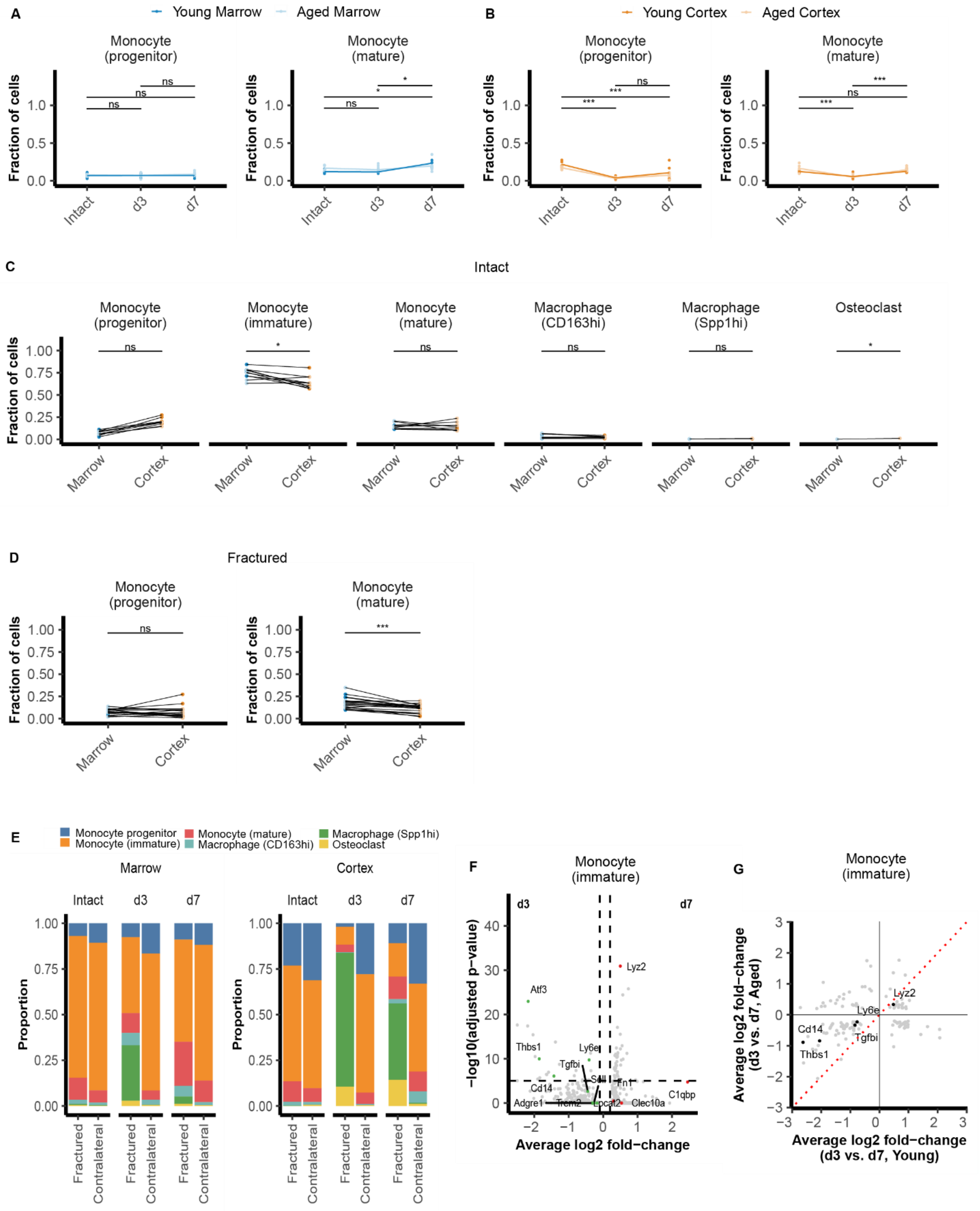
A
Young Marrow
Aged Marrow
B
Young Cortex
Aged Cortex
Monocyte (progenitor)
Monocyte (mature)
Fraction of cells
Intact
d3
d7
C
Intact
Monocyte (immature)
Macrophage (CD163hi)
Macrophage (Spp1hi)
Osteoclast
Marrow
Cortex
D
Fractured
E
Monocyte progenitor
Monocyte (immature)
Monocyte (mature)
Macrophage (CD163hi)
Macrophage (Spp1hi)
Osteoclast
Proportion
Fractured
Contralateral
F
−log10(adjusted p−value)
Average log2 fold−change
Lyz2
Atf3
Thbs1
Ly6e
Tgfbi
Sell
Cd14
Fn1
C1qbp
Adgre1
Trem2
Lpcat2
Clec10a
G
Average log2 fold−change (d3 vs. d7, Aged)
Average log2 fold−change (d3 vs. d7, Young)

**Suppl. Fig. 5: Distinct temporal and tissue-specific differentiation trajectories of monocyte-derived macrophages during early fracture healing.**

(A-B) Fractional abundance of monocyte progenitors and mature monocytes across intact, day 3, and day 7 conditions in young (blue) and aged (orange) mice, showing distinct dynamics over time.

(C-D) Paired comparisons of monocyte and macrophage subtypes between marrow and cortex within replicates in (C) intact samples and (D) fractured samples.

(E) Stacked bar plots showing proportional distribution of monocyte/macrophage subsets across marrow and cortex compartments at all timepoints, comparing fractured and contralateral sides.

(F) Volcano plot showing differentially expressed genes in immature monocytes at day 3 vs. day 7 post-osteotomy in young mice, highlighting changes associated with early activation and remodeling.

(G) Comparative gene expression plot showing $log_2$ fold-change in immature monocytes between day 3 and day 7 in young versus aged mice. Red dotted line indicates identity (no age effect).

n = 4-5 technical replicates per group; *p < 0.05, **p < 0.01, ***p<0.001; ns = not significant, Wilcoxon test or paired test with Benjamini-Hochberg correction for multiple testing.
